# Dynamic trajectory cues drive sequenced integration in approach detectors

**DOI:** 10.64898/2026.06.05.730527

**Authors:** Harsh Vashistha, Natalia CB matos, Heng Wu, Damon Clark

**Affiliations:** Department of Molecular, Cellular and Developmental Biology, Yale University; Quantitative Biology Institute, Yale University; Interdepartmental Neuroscience Program, Yale University; Department of Physics, Yale University; Department of Neuroscience, Yale University; Wu Tsai Institute, Yale University

## Abstract

Detecting approaching objects is essential for survival, and studies across animals have identified neurons and circuits selective for objects that grow in size over time, a hallmark of visual approach. However, approach as a physical process generates a rich ensemble of correlated cues governed by geometry and physics, of which size expansion is just one, and how the brain incorporates this broader cue structure to detect approach remains unknown. Here, we show that a visual stimulus without size expansion but with modulated luminance, a cue intrinsic to approach, elicits compelling percepts of approach and retreat in both humans and the fruit fly Drosophila, revealing an unsuspected, cross-species percept of approach based on luminance alone. Using targeted genetic silencing and two-photon calcium imaging in flies, we identify neurons that mediate evasive responses to both expansion-based and luminance-based approach, establishing them as general approach detectors. Guided by the geometry of approach, we show that luminance cues should precede detectable expansion signals during naturalistic approach. Consistent with this, these neurons combine luminance and expansion cues synergistically, but only when luminance changes precede expansion, matching their natural temporal order, thus performing sequenced cue integration. Together, these findings anchor a cross-species perceptual phenomenon to a defined circuit computation and reveal how neural circuits are tuned to the dynamic cue structure of natural events.

## INTRODUCTION

Detecting approaching objects, such as predators or objects on a collision course, is a fundamental visual task for sighted animals. Such threats trigger rapid defensive or evasive behaviors across phyla (Ball and Tronick, 1971; Carbone et al., 2018; Gabbiani et al., 2002; Heinemans and Moita, 2024; Liu et al., 2011; Schiff et al., 1962; Temizer et al., 2015; Yilmaz and Meister, 2013; Zhao et al., 2014) including freezing (Ferreira and Moita, 2020; Franceschi et al., 2016), turning (Kim et al., 2023; Muijres et al., 2014; Tammero and Dickinson, 2002), escape(Ache et al., 2019; Dunn et al., 2016; Evans et al., 2018; Ishikane et al., 2005; Oliva and Tomsic, 2012; Reyn et al., 2017), or postural adjustments (Card and Dickinson, 2008; Dombrovski et al., 2023), all to minimize predation and collision risk.

Visual approach detection is fundamentally an inference problem in which the brain must estimate decreasing object distance from inherently ambiguous retinal cues. Several visual cues that signal relative distance, including looming, occlusion, luminance and contrast, could, in principle, contribute to approach detection (Coules, 1955; Cutting and Vishton, 1995; Farne, 1977; O’Shea et al., 1994; Sun et al., 2016). Among these, looming — the progressive increase in angular size — is the most salient and has historically dominated studies of approach detection (Ache et al., 2019; Clarke et al., 2013; Dunn et al., 2016; Fotowat and Engert, 2023; Gabbiani et al., 2002; Klapoetke et al., 2017; Liu et al., 2011; Peron and Gabbiani, 2009; Schiff et al., 1962; Sun and Frost, 1998; Temizer et al., 2015; Wang and Frost, 1992; Yamawaki and Toh, 2009; Zhou et al., 2022). However, a single cue such as expansion may be in-sufficient for reliably detecting approach in all contexts. At long distances, an object’s apparent size may fall below the spatial resolution of the visual system, and geometry constrains its rate of angular expansion, rendering it potentially undetectable by motion sensors. Similarly, even when expansion is detectable, visual noise can degrade expansion signals, making additional cues valuable for inferring approach. In such conditions, where expansion signals are weak or absent, additional cues, such as luminance change, could provide complementary information about approach.

Psychophysical studies in humans have consistently shown that manipulations of visual brightness or contrast can alter perception of object distance (Ashley, 1898; Egusa, 1983; Farne, 1977; Hibbard et al., 2023; O’Shea et al., 1994; Schwartz and Sperling, 1983): brighter stimuli and higher-contrast targets tend to appear nearer, particularly when other depth cues are limited or absent. This perceptual bias likely reflects an ecological regularity: in natural scenes, luminance has a modest negative correlation with distance due to ambient occlusion, such that more distant surfaces receive less diffuse illumination and appear darker (Cooper and Norcia, 2014; Lee and Potetz, 2003; Samonds et al., 2012; Scaccia and Langer, 2018).This static bias has a dynamic counterpart: as an object approaches an observer, it not only grows in angular size but also progressively occludes the background, replacing it with its own luminance, producing a change in the overall luminance of the visual field. Together these observations raise the possibility that the nervous system exploits natural regularities to integrate luminance and expansion cues and improve detection. Yet how luminance change contributes to approach detection, and how it is integrated with expansion in identified circuits, remain unknown.

Here, we first employ a purely luminance-based stimulus, devoid of expansion, to isolate the role of luminance change in approach detection. We combine human psychophysics with fly behavioral measurements to demonstrate that luminance change serves as a complementary cue for detecting approach and retreat in the absence of expansion. Using two-photon imaging and genetic perturbations in the fly, we identify neural loci where luminance and expansion cues are combined in the brain. There, we uncover a sequenced cue integration mechanism that combines these two cues to generate an amplified approach signal. Together, these results define the circuit basis for a purely luminance-driven approach percept shared between flies and humans, and reveal how natural temporal cue structure shapes circuit computations to produce robust signals that guide evasive behaviors critical for survival.

## RESULTS

### A luminance-based stimulus generates strong approach and retreat percepts in humans

To isolate the contribution of luminance from that of expansion in approach detection, we sought a visual stimulus that can evoke a strong percept of approach without changing object size. Earlier work had shown that a grid of dots with changing luminance could elicit robust depth-motion percepts in humans while keeping stimulus size constant (Weiss et al., 2004). We adapted this stimulus and created a 6 × 6 grid of circles, which we term a luminance-grid stimulus (see Methods, Supplemental Video S1). The luminance value of each circle followed either an increasing sawtooth waveform (ramp-up) or a decreasing sawtooth waveform (ramp-down) with a period of 0.5 s (Figure 1a). The initial luminance values of each circle were drawn uniformly from between 0 and 1, ensuring that the intensity across all circles averaged to gray (0.5) and that on average no net directional motion was present in the stimulus.

**Figure 1:**
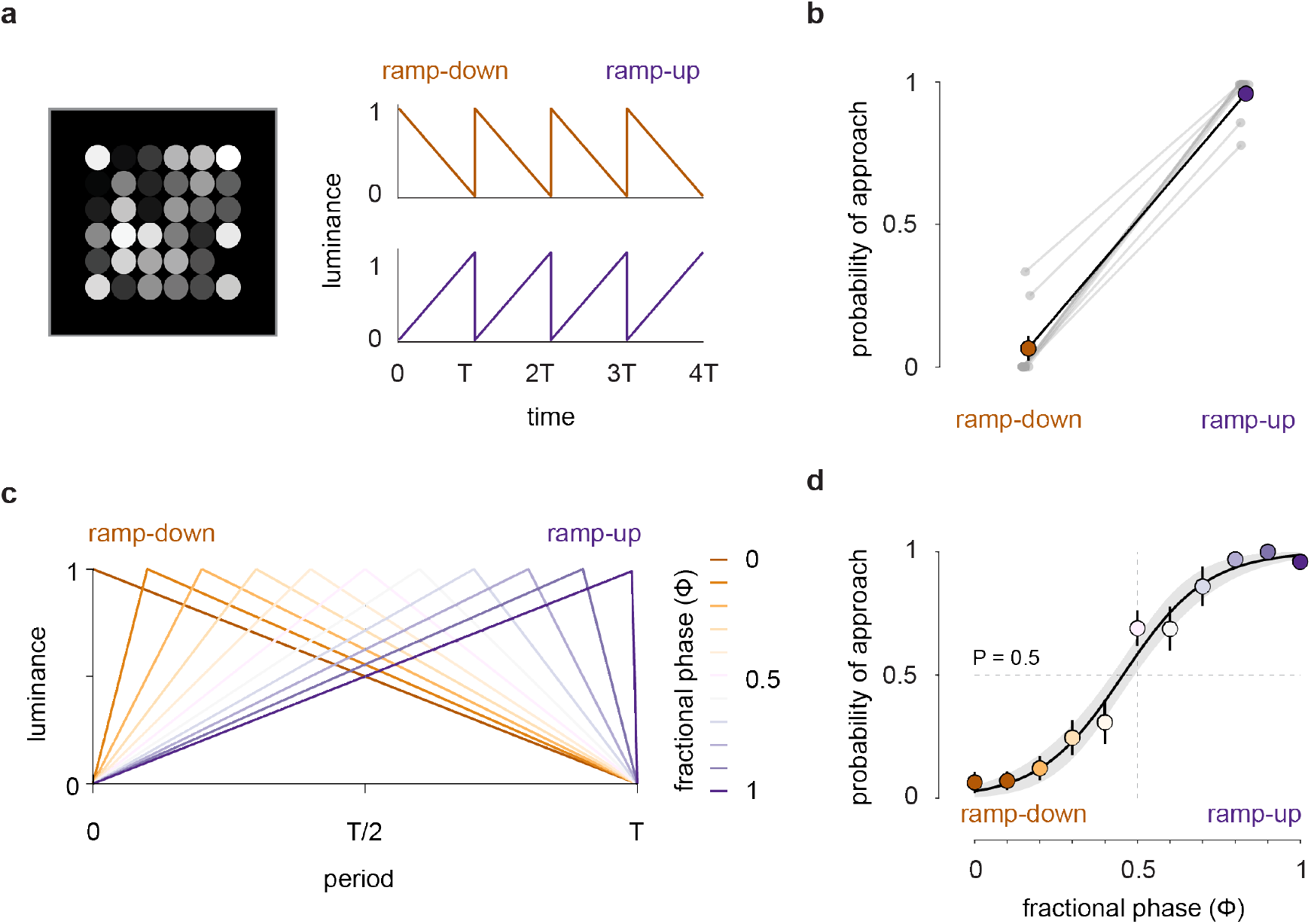
Humans perceive approach and retreat in a purely luminance-based stimulus. **(a)** A purely luminance-based stimulus evokes perception of approach and retreat in human subjects (Supplemental Video 1). The luminance of each circle is modulated using either a ramp-up or ramp-down sawtooth waveform with a 0.5 s period. **(b)** Probability of approach perception in human subjects (n = 9). The ramp-up stimulus evoked a strong perception of approach, while the ramp-down stimulus evoked a strong perception of retreat. Solid error bars represent the standard error of the mean (SEM). Linked gray dots are individual subjects. **(c)** Interpolated version of the original stimulus time trace parameterized by fractional phase (*ϕ*), representing the fraction of ramp-up duration within one period. **(d)** Probability of approach perception as a function of fractional phase (*ϕ*) of the stimulus. Approach perception increased monotonically, following a sigmoidal function. Solid error bars represent the standard error of the mean (SEM), and the shaded area represents the 95% functional bounds of the best fit.

We presented these ramp-up and ramp-down stimuli to human subjects and performed a two-alternative forced-choice (2AFC) task to quantify the probability of reporting approach in each condition. Ramp-up stimuli reliably evoked strong illusory percepts of approach, while ramp-down stimuli evoked equally strong percepts of retreat in human subjects, consistent with earlier findings (Weiss et al., 2004) (Figure 1b, Figure S1a). These results confirm that luminance change alone is sufficient to drive approach and retreat percepts in the absence of expansion.

Because the ramp-up and ramp-down stimuli differ in their luminance asymmetry, with ramp-up producing slow luminance increments followed by fast decrements and ramp-down producing slow luminance decrements followed by fast increments, we asked how the balance of luminance asymmetry drives approach and retreat percepts. We designed a family of luminance waveforms by varying the relative durations of the slow-increase (ramp-up) and slow-decrease (ramp-down) phases within a cycle (Figure 1c), parameterized by a fractional phase (Φ), representing the proportion of the cycle devoted to slow luminance increase. Thus, Φ = 0 corresponded to pure ramp-down, Φ = 1 to pure ramp-up, and Φ = 0.5 to a symmetric triangular wave with equal increasing and decreasing phases (Supplemental Video S2). We repeated our 2AFC task across this continuum of luminance-grid stimuli. The probability of approach perception increased monotonically with Φ, following a sigmoidal function (Figure 1d). At Φ = 0.5, the probability of perceiving approach was near chance, indicating that balanced increments and decrements in luminance largely abolished approach and retreat percepts. This is consistent with the notion that approach and retreat percepts in this stimulus emerge from temporal asymmetry in the luminance increments and decrements.

However, experiments with human subjects cannot easily establish the mechanistic origin of these percepts or identify the neural substrates that generate them. We therefore turned to the fruit fly *Drosophila*, a genetically tractable model with well-mapped visual circuitry, to identify neural bases of luminance-based approach detection.

### Fly responses to luminance-grid stimuli mirror human reports

Fruit flies display robust defensive behaviors, including freezing and evasive turning, when exposed to expanding (looming) stimuli simulating approaching objects (Ache et al., 2019; Card and Dickinson, 2008; Ferreira and Moita, 2020; Muijres et al., 2014; Reyn et al., 2017; Zacarias et al., 2018). The freezing response, characterized by a rapid cessation of walking, is thought to represent a “freeze-and-assess” strategy that reduces visibility to predators (Ferreira and Moita, 2020), while directed turns away from a looming stimulus during flight facilitate evasion (Muijres et al., 2014). We asked whether luminance-grid stimuli could drive similar freezing and turning behaviors in walking flies, which would strongly suggest that these stimuli generate approach and retreat percepts in flies as they do in humans.

Using a custom fly psychophysics setup (Creamer et al., 2019) (Figure 2a), we recorded behavioral responses of walking flies to luminance-grid stimuli. We first established baseline responses to classical size-based approach and retreat signals by presenting expanding and contracting disc stimuli at 60°/s in our tethered walking preparation (Figure 2b). Stimuli were presented on ±90° azimuthally relative to the fly’s heading and directly ahead of the fly while turning and freezing were recorded.

**Figure 2:**
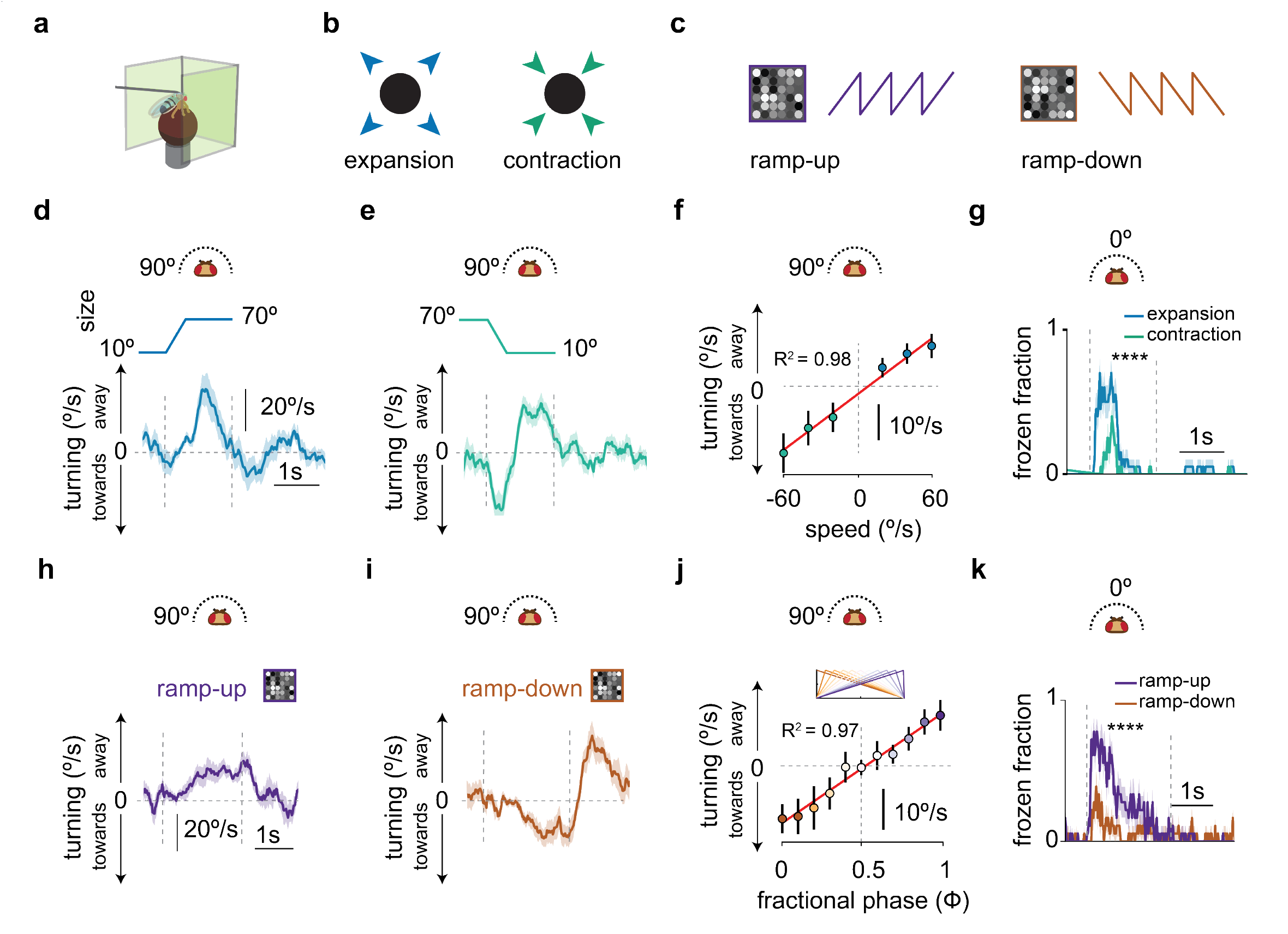
Fly responses to luminance-grid stimuli mirror human reports. **(a)** Fly-on-a-ball setup to measure fly behavioral responses to visual stimuli. **(b)** Expanding and contracting disc stimuli. **(c)** Ramp-up and ramp-down luminance grid stimuli. **(d)** Turning response of flies to a disc expanding at 60°/s, presented at 90° azimuthally relative to the fly’s heading. **(e)** Turning behavior of flies to a disc stimulus contracting at 60°/s, presented at 90° azimuthally. **(f)** Mean turning response of flies to disc stimuli presented at 90° azimuthally, contracting or expanding at −60°/s to 60°/s. Mean turning was calculated from onset to the end of contraction and from onset to 0.75 s after reaching the final size for expansion. Red solid line represents the linear best fit (*R*^2^ = 0.98, p = 0.001). **(g)** Freezing response of flies to a disc stimulus presented in front, either expanding or contracting at 60°/s (p < 0.0001). **(h)** Turning response of flies to a ramp-up luminance-grid stimulus presented at 90° azimuthally relative to the fly’s heading (period = 0.5 s). **(i)** Turning response of flies to a ramp-down luminance-grid stimulus presented at 90° azimuthally relative to the fly’s heading (period = 0.5 s). **(j)** Mean turning response of flies to ramp-up and ramp-down stimuli. Mean turning responses were calculated from onset to the end of stimulus presentation. Red solid line represents the linear best fit (*R*^2^ = 0.97, p < 10^−5^). **(k)** Freezing response of flies to luminance-grid stimuli presented in front of fly (p < 0.0001). Vertical dotted lines mark the onset of expansion or contraction and the bounds of the luminance-grid stimulus. Shaded area represents standard error of the mean (SEM). n = 20 flies for (d–g) and n = 17 flies for (h–k). p-values were calculated using Student’s t-test (**** p < 0.0001, *** p < 0.001, ** p < 0.01, * p < 0.05).

Walking flies exhibited slow ramped turns away from the expanding disc stimuli presented laterally (Figure 2d), demonstrating robust looming-evoked evasive responses in walking flies. Although evasive turning has previously been reported in flying flies (Kim et al., 2023; Muijres et al., 2014), these results establish that evasive turning is not flight specific. In contrast, contracting discs evoked an opposite turning response, with flies initially turning towards the stimulus during its contraction, followed by turning away from it as it persisted as a small dot (Figure 2e). The strength of the turning response to expanding and contracting discs depended on the expansion and contraction speed (Figure 2f), confirming that flies are sensitive to the rate of size change as an approach cue.

When expanding and contracting disc stimuli were presented directly in front of the fly, expansion elicited a strong, transient freezing response, while contraction elicited only minimal freezing (Figure 2g, Figure S1b). These results demonstrate that walking flies exhibit location- and stimulus-specific defensive responses where expansion drives evasive turning and freezing, while contraction drives approach turning and elicits minimal freezing.

We next presented flies with ramp-up and ramp-down luminance-grid stimuli (Figure 2c) and compared their responses to those evoked by expanding and contracting discs. The ramp-up stimulus drove strong turns away from the stimulus (Figure 2h) and robust freezing (Figure 2k), mirroring responses to expanding discs, while ramp-down stimulus drove turns towards the stimulus (Figure 2i) and minimal freezing (Figure 2k), mirroring responses to contracting discs. These behavior results parallel the human judgements of approach and retreat observed in our psychophysical assay (Figure 1b), suggesting that flies may share the same percepts of these luminance-based stimuli.

To draw the comparison further, we recorded fly responses to the same continuum of interpolated luminance-grid stimuli presented to humans (Figures 1d and 2j). Fly turning responses shifted from turning toward to away as the stimulus progressed from ramp-down to ramp-up, closely matching the human psychophysical curve for approach perception. As in humans, at Φ = 0.5, where luminance increments and decrements are balanced, flies exhibited little net turning. While these behavioral results in flies are suggestive, it is in principle possible that they emerge from fly preferences for distinct spatiotemporal patterns, unrelated to approach percepts.

### Silencing loom-detectors suppresses fly behavioral response to luminance-grid stimuli

To better link the luminance-grid responses to percepts of approach, we asked which neural circuits mediate the luminance grid responses. Several neural populations have been identified in the fly brain as necessary for defensive responses to expanding stimuli (Ache et al., 2019; Kim et al., 2023; Klapoetke et al., 2017, 2022; Reyn et al., 2017; Vries and Clandinin, 2012; Wu et al., 2016), including lobula columnar (LC) and lobula plate–lobula columnar (LPLC) neurons. These neurons are characterized as loom detectors based on their responses to stimuli that increase in size. However, in natural contexts, approaching objects produce multiple co-varying visual cues including retinal expansion and systematic luminance changes. It is therefore unclear whether these neurons detect approaching objects based on expansion alone or whether they integrate multiple correlated visual cues to signal approach more generally.

We hypothesized that neurons classically associated with loom-detection might encode a general approach signal rather than one exclusively tied to expansion. If so, these neurons could mediate behavioral responses both to classical expanding stimuli and to luminance-grid approach stimuli, and silencing them would disrupt responses to both stimulus classes. Disruption of responses to luminance-grid stimuli upon silencing would therefore identify the circuits mediating those responses. Crucially, such disruption would also support the interpretation that fly behavioral responses to luminance-grid stimuli reflect percepts of approach or retreat rather than arbitrary spatiotemporal preferences.

To test this hypothesis, we used the Gal4–UAS system to express a temperature-sensitive suppressor of synaptic transmission, *shibire*^ts^ (Kitamoto, 2001), to reversibly silence two well-characterized loom-selective cell types, LPLC1 and LPLC2 (Ache et al., 2019; Kim et al., 2023; Klapoetke et al., 2017, 2022; Tanaka and Clark, 2022; Wu et al., 2016) (Figure 3a, Figure 3b). These neurons mediate behavioral responses to looming stimuli, including escape take-off (Ache et al., 2019; Reyn et al., 2017) and turning responses during flight (Kim et al., 2023). As a control, we silenced LC11, a cell type that does not respond to looming but mediates freezing responses to small moving objects (Keleş and Frye, 2017; Keleş et al., 2020; Tanaka and Clark, 2020). With these neurons silenced, we quantified freezing and turning responses in our tethered walking assay across four stimulus conditions: expanding and contracting discs and ramp-up and -down luminance-grid stimuli.

**Figure 3:**
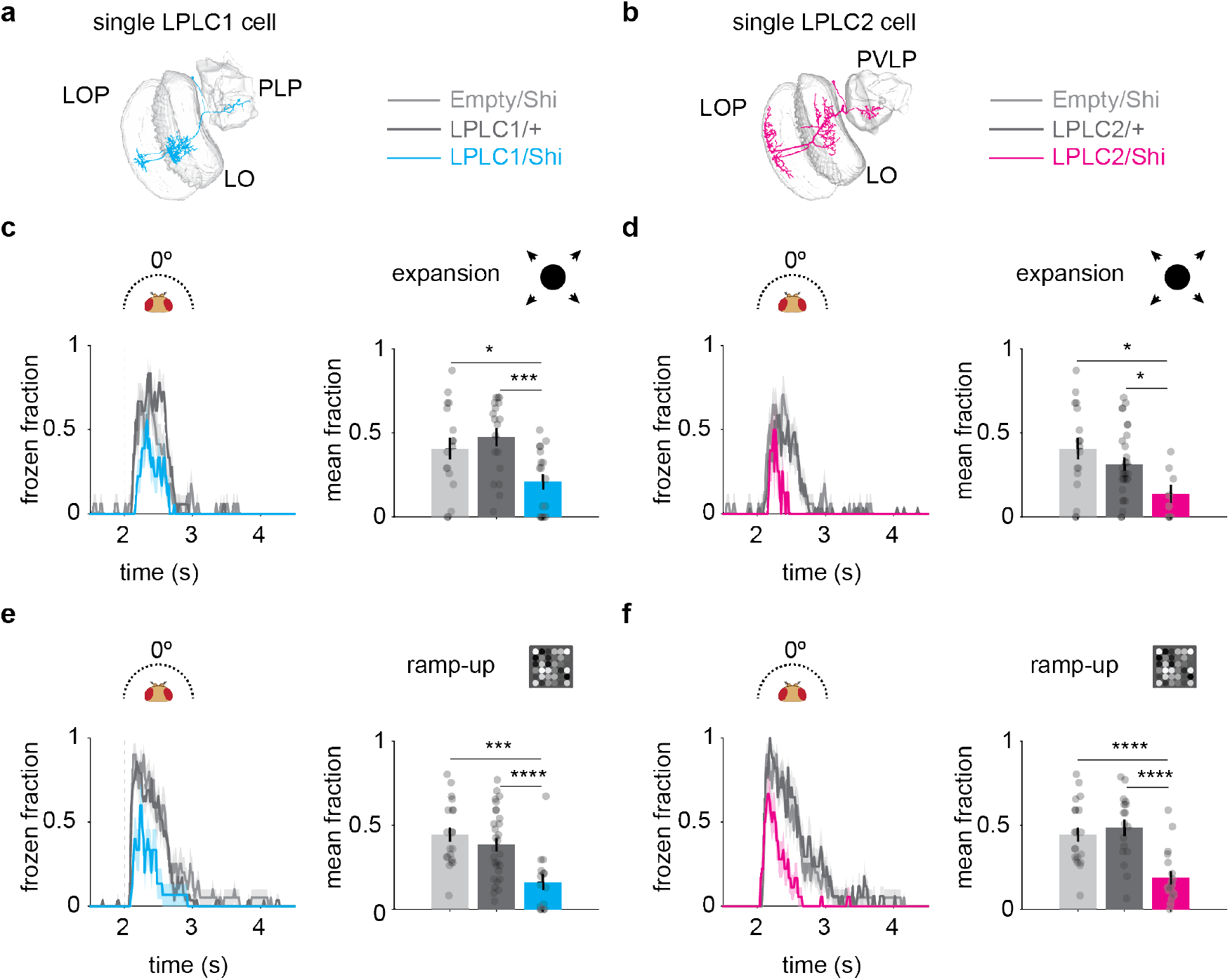
LPLC1 and LPLC2 neurons mediate freezing responses to approach. **(a–b)** Location of LPLC1 and LPLC2 neurons in the fly brain. LPLC1 and LPLC2 neurons receive inputs in the lobula (LO) and lobula plate (LOP) regions and project to the posterior lateral protocerebrum (PLP) and posterior ventrolateral protocerebrum (PVLP) regions, respectively. **(c)** Silencing LPLC1 neurons significantly reduced the freezing response to an expanding disc stimulus (60°/s) in comparison to controls. Empty/Shi (n = 17 flies), LPLC1/+ (n = 18 flies), LPLC1/Shi (n = 18 flies). **(d)** Silencing LPLC2 neurons significantly reduced the freezing response to an expanding disc stimulus (60°/s) in comparison to controls. Empty/Shi (n = 17 flies), LPLC2/+ (n = 18 flies), LPLC2/Shi (n = 8 flies). **(e)** Silencing LPLC1 neurons significantly reduced the freezing response to a ramp-up stimulus (period = 0.5 s) in comparison to controls. Empty/Shi (n = 20 flies), LPLC1/+ (n = 29 flies), LPLC1/Shi (n = 15 flies). **(f)** Silencing LPLC2 neurons significantly reduced the freezing response to a ramp-up stimulus (period = 0.5 s) in comparison to controls. Empty/Shi (n = 20 flies), LPLC2/+ (n = 17 flies), LPLC2/Shi (n = 18 flies). Mean frozen fraction was calculated in a 1-s window starting at the stimulus onset in all conditions. Gray dots are individual flies. Vertical dotted lines mark the onset of expansion and luminance-grid modulation. Shaded area represents standard error of the mean (SEM). Stars show significance level; p-values were calculated using Student’s t-test (**** p < 0.0001, *** p < 0.001, ** p < 0.01, * p < 0.05).

Silencing LPLC1 and LPLC2 neurons significantly reduced freezing responses to the expanding disc (Figure 3c, Figure 3d), consistent with prior silencing experiments and measurements showing loom sensitivity (Ache et al., 2019; Kim et al., 2023; Klapoetke et al., 2017, 2022). Interestingly, we also observed deficits in the freezing response to ramp-up stimuli when each cell type was silenced, relative to genetic controls (Figure 3e, Figure 3f). These deficits demonstrate that LPLC1 and LPLC2 neurons mediate freezing responses to both size-based and luminance-based stimuli. These results further suggest that luminance-grid stimuli evoke a genuine percept of approach in flies mediated by loom-sensors. Silencing LC11 neurons did not affect either turning or freezing responses to any stimulus (Figure S1, Figure S2). These findings support the notion that LPLC1 and LPLC2 neurons encode a general approach signal integrating both luminance and expansion cues, rather than functioning as simple expansion detectors.

Analysis of turning behavior revealed that silencing LPLC1 neurons significantly reduced turning away from ramp-up stimuli and, surprisingly, enhanced turning toward ramp-down stimuli (Figure S3c, Figure S3g) and contracting discs (Figure S3e). In contrast, silencing LPLC2 selectively impaired turning away from ramp-up stimuli (Figure S3d) without significantly affecting turning toward ramp-down stimuli (Figure S3h).

Together, these results indicate that, like humans, flies perceive approach and retreat in luminance-grid stimuli, recruiting established looming-selective circuits to guide behavior. The fact that flies and humans share these percepts in motionless stimuli suggests an evolutionarily widespread computational strategy for inferring approach and retreat through the integration of multiple visual cues, including both luminance change and expansion.

### LPLC1 and LPLC2 respond differentially to ramp-up and ramp-down stimuli

To better understand how LPLC1 and LPLC2 neurons encode approach and retreat signals, we expressed the calcium indicator GCaMP6f (Chen et al., 2013) in each cell type and used two-photon calcium imaging (Figure 4a) to measure responses to the four stimuli defined previously (Figures 4b and 4c): expanding and contracting discs and ramp-up and ramp-down luminance-grid stimuli.

**Figure 4:**
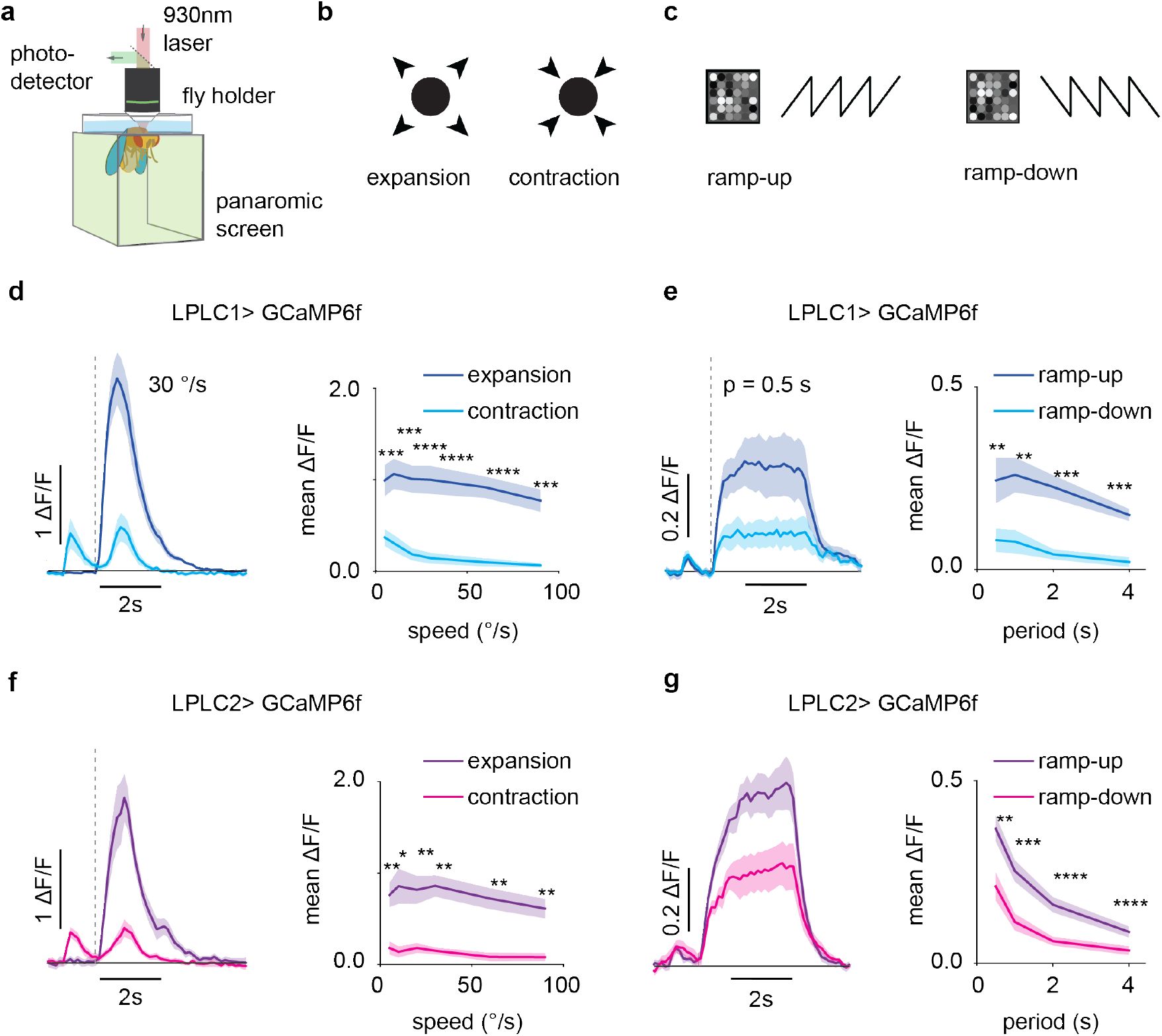
LPLC1 and LPLC2 neurons respond differentially to expansion and contraction stimuli and to ramp-up and ramp-down luminance-grid stimuli. **(a)** Two-photon imaging setup used for in vivo calcium recordings during visual stimulation. **(b–c)** Four stimulus classes used for visual stimulation: expanding disc, contracting disc, and ramp-up and ramp-down luminance-grid stimuli. **(d)** Calcium responses in LPLC1 neurons (n = 8 flies) to expanding and contracting disc stimuli for a range of radial speeds (5°/s, 10°/s, 20°/s, 30°/s, 60°/s, 90°/s). **(e)** Calcium responses in LPLC1 neurons (n = 7 flies) to ramp-up and ramp-down stimuli for a range of sawtooth periods (period = 0.5 s, 1 s, 2 s, 4 s). **(f)** As in (d) but for LPLC2 neurons (n = 6 flies). **(g)** As in (e) but for LPLC2 neurons (n = 8 flies). Vertical dotted lines mark the onset of expansion (contraction) and luminance-grid modulation. Shaded areas represent the standard error of the mean (SEM). Stars show significance level; p-values were calculated using a paired t-test (**** p < 0.0001, *** p < 0.001, ** p < 0.01, * p < 0.05). See Methods for detailed stimulus description.

We first presented linearly expanding and contracting dark discs at six speeds, centered on the receptive fields of these neurons (see Methods). Both LPLC1 and LPLC2 neurons showed strong selectivity for expanding over contracting discs at all speeds (Figure 4d, Figure 4f), consistent with their well-established sensitivity to looming stimuli (Ache et al., 2019; Klapoetke et al., 2017, 2022; Tanaka and Clark, 2022).

We next asked whether these neurons respond differentially to the ramp-up and ramp-down luminance-grid stimuli. We presented ramp-up and ramp-down stimuli with four sawtooth wave periods. Because these stimuli do not change in size and span a large visual area, we measured calcium responses in the axon terminals of these neurons in their optic glomeruli. Both LPLC1 and LPLC2 neurons responded more strongly to the ramp-up than to the ramp-down stimuli across all periods (Figure 4e, Figure 4g, Figure S4). This differential sensitivity mirrors the behavioral asymmetry in ramp-up and ramp-down stimuli and is consistent with their role in approach-selective defensive behavior. Together, these imaging results show that LPLC1 and LPLC2 neurons are tuned to both size-based and luminance-based cues, supporting their roles as general approach detectors that drive defensive behavior (Figure 2, Figure 3).

### Luminance cues precede expansion cues in visual systems with limited resolution

We next asked how luminance and expansion cues evolve during naturalistic approach events. As an object approaches, the sensory signals available to an observer evolve over time based on the geometry of approaching objects (Figure 5). For a distant object approaching at constant velocity, the apparent size of the object increases non-linearly on the retina (Gabbiani et al., 2002; Lee, 1976) (Figure 5b). At long distances, the apparent size (*θ*) scales with distance as 1/*D*, where *D* is the distance between the object and the observer. However, at those same distances, the irradiance at the eyes due to the object’s luminance scales as 1/*D*^2^. This means that when the object is moving toward the viewer, the time derivative of irradiance changes more rapidly (1/*D*^3^) than of angular size (1/*D*^2^) (see Appendix). However, since expansion signals may be measured along the object’s entire perimeter, which scales like 1/*D*, both the irradiance derivative and expansion signal scale with distance as 1/*D*^3^ (see Appendix). Thus, both signals increase proportionally as an object approaches.

**Figure 5:**
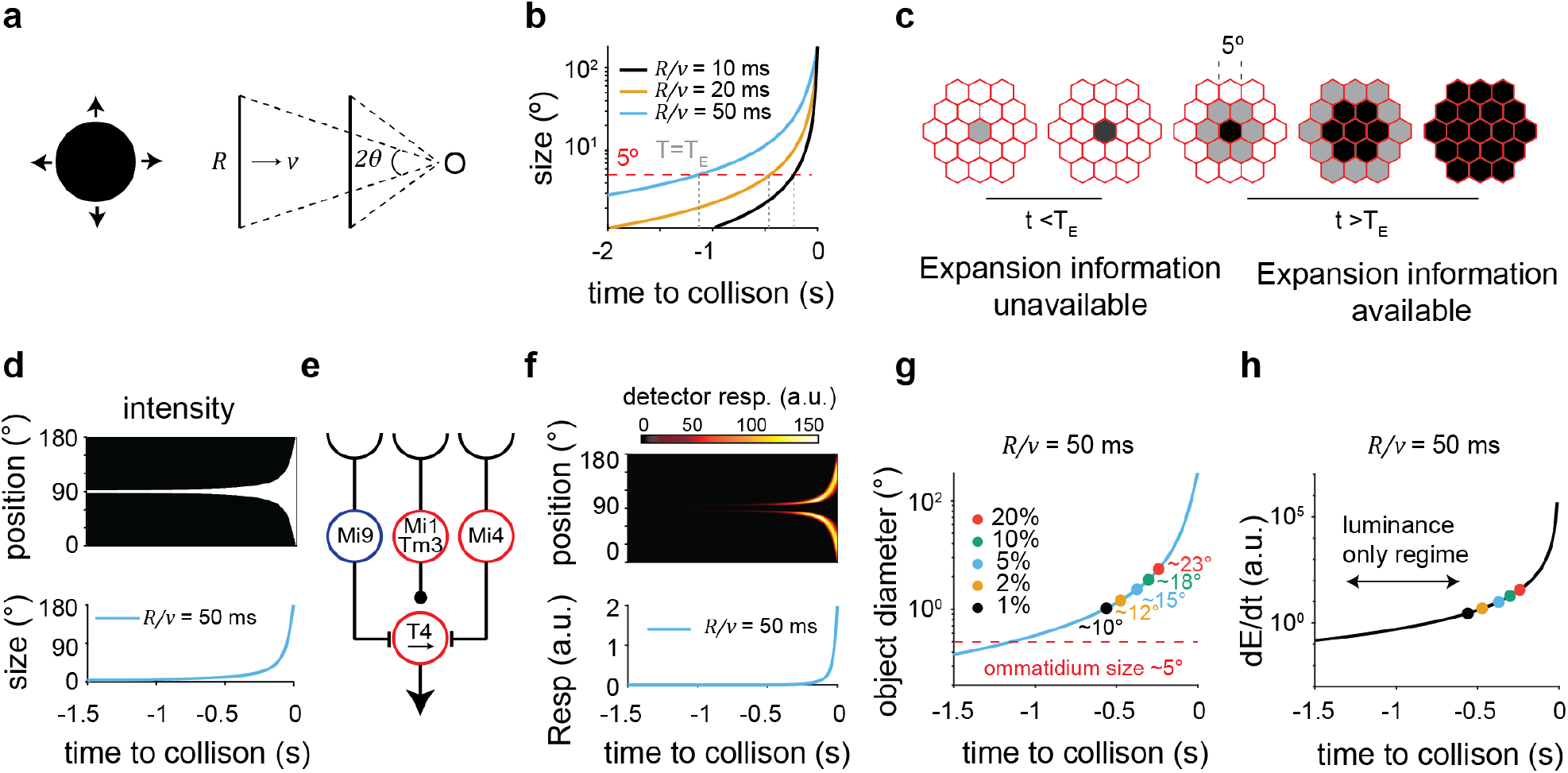
Luminance cues precede expansion signals during approach. **(a)** Illustration of the approach dynamics of an object of size *R* approaching with speed *v*. **(b)** Temporal dynamics of angular size as the object approaches the observer with different *R*/*v* ratios. **(c)** Illustration of the availability of expansion and luminance information in two temporal regimes based on the spatial resolution limit of the flies. **(d)** One-dimensional representation of looming intensity with *R*/*v* = 50 ms. **(e)** A minimal synaptic model for light-edge-selective T4 neurons. **(f)** Top panel: simulated calcium proxy response of T4-like detectors to a one-dimensional expanding edge (as in (d)). Bottom panel: estimated response of a population of motion detectors to two-dimensional expanding stimuli as a function of time, obtained by summing the one-dimensional radial calcium proxy across space (*x* ∈ [0, *θ*(*t*)]) and scaling by 2*πθ*(*t*) to account for the disc circumference. **(g)** Object diameter at which motion signals cross different thresholds (1%–20% of the response of the motion detectors at an object diameter of 40° in this simulation; see Methods). **(h)** Rate of change of irradiance (Appendix 1) due to an object approaching at *R*/*v* = 50 ms. Colored dots mark the time to collision at different motion signal thresholds, as in (g).

Importantly, however, when an object is far away, its angular subtense may fall below the spatial resolution threshold of the visual system (typically considered ∼5° for flies (Stavenga, 2003)) (Figure 5c). When the object is so small, its radial expansion is likely unresolvable by motion detectors that sense motion across multiple ommatidia (Shinomiya et al., 2019; Takemura et al., 2017), making expansion signals potentially unobtainable. Yet, in these conditions the integrated luminance within the visual field may still change detectably, providing an early signal of approach before expansion becomes resolvable (Figure 5c). As the object approaches, its angular size and velocity eventually become detectable by motion detectors. Before that time, changes in local average luminance can in principle signal approaching stimuli.

To gain intuition about these two detection mechanisms, we simulated an approaching object projected onto the retina (Figure 5d) while using a minimal synaptic model (Zavatone-Veth et al., 2020) (Figure 5e, Figure 5f) to estimate associated motion signals from elementary motion detectors T4 and T5. We then asked when, with respect to collision time, motion signals rose above different thresholds, mimicking a form of expansion detection threshold. Depending on the threshold, detection of expansion can occur when the object is substantially larger than one ommatidium (Figure 5g). However, the luminance signal remains available and informative to the observer during all portions of the approach, even when expansion is unresolvable (Figure 5h). That is, an integrated luminance cue can be useful even when integrating over regions larger than a single ommatidium.

### Natural sequencing enhances responses to combined luminance and expansion stimuli

Based on the intuition gained by the simulation, we concluded that the temporal sequence of local luminance change preceding an expansion signal constitutes a reliable naturalistic signature of object approach. We hypothesized that LPLC1 and LPLC2 could be specifically tuned to exploit this ordering for early approach detection. To test this hypothesis, we first examined the responses of LPLC1 and LPLC2 neurons to isoluminant expansion and to luminance changes in isolation. We designed an isoluminant expanding annulus with an inner radius of 5° and an outer radius of 15°. The annulus contained a checkerboard pattern with a check size of 5° × 22.5° (radial coordinate in visual angle × annulus angular coordinate), ensuring that average luminance changes during expansion were eliminated (Figure 6a; Supplemental Video S3). The annulus expanded to final inner and outer radii of 20° and 30°, moving at 30°/s. In natural scenes, approaching objects can produce local luminance changes that are either increases or decreases from the background, depending on whether the object is lighter or darker than the background. Therefore, to isolate luminance-change responses, we presented a gray-to-white or gray-to-black dot transition at the receptive-field centers of the neurons of interest (Figures 6b and 6c, Figure S5).

**Figure 6:**
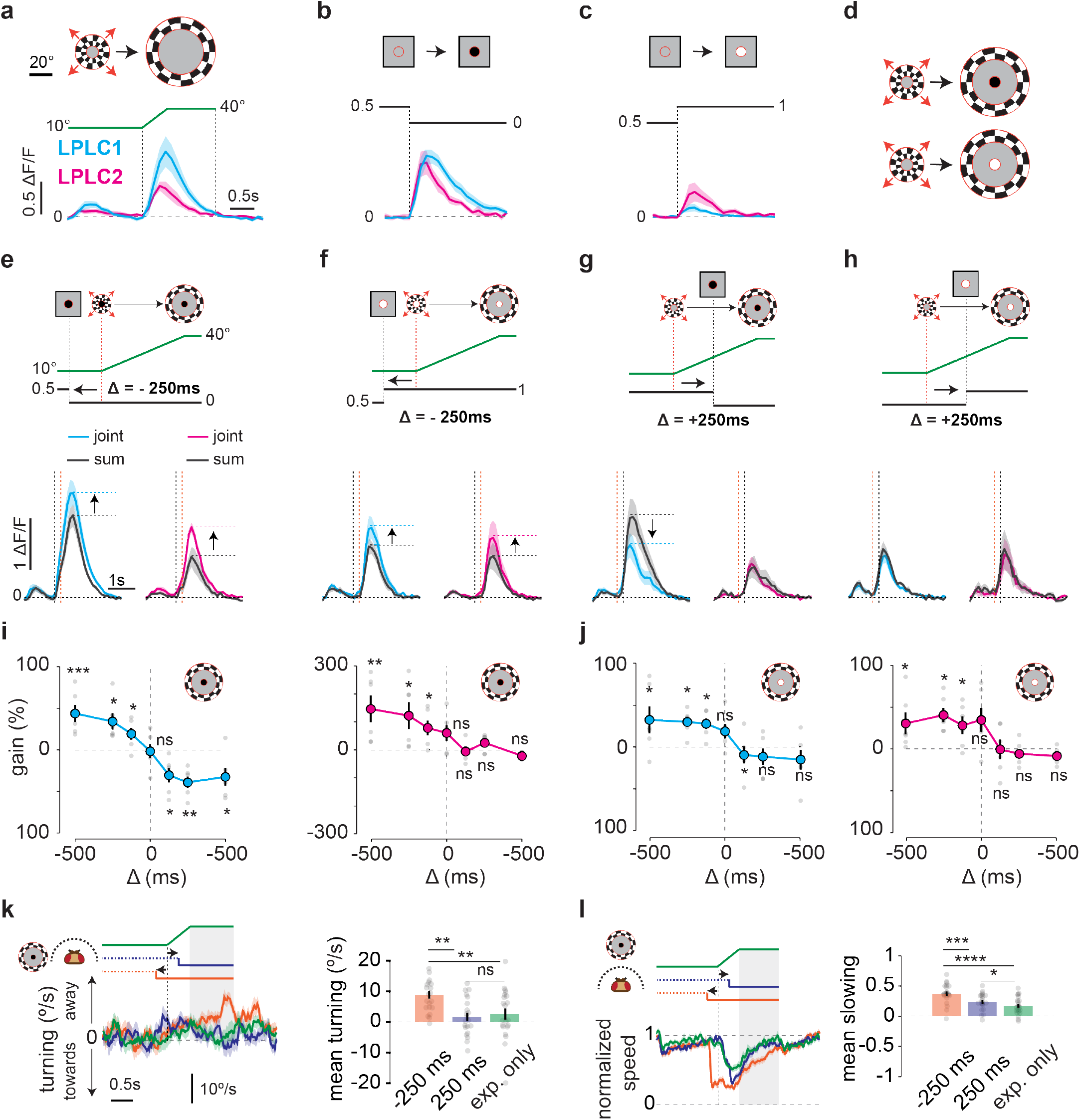
LPLC1 and LPLC2 neurons sequentially integrate luminance and expansion cues. **(a)** Responses of LPLC1 and LPLC2 neurons to isolated presentation of an isoluminant annulus. The annulus had an initial inner radius of 5°, an outer radius of 15°, and a checkerboard pattern with a check size of 5° × 22.5° (radial coordinate in visual angle × annulus angular coordinate). The annulus inner and outer radii each expanded at 30°/s until the outer radius reached 30°. **(b)** Responses of LPLC1 (n = 6 flies) and LPLC2 (n = 6 flies) to a black dot (radius 5°) presented at the receptive-field (RF) center to introduce an isolated luminance change. **(c)** As in (b) but with a white dot. **(d)** Joint stimuli in which the expanding annulus and the luminance change were combined at different temporal offsets. **(e)** Calcium responses of LPLC1 (n = 6 flies) and LPLC2 (n = 6 flies) neurons to the joint presentation of isoluminant expansion and a gray-to-black luminance change at a time offset of Δ = −250 ms, compared to the expected linear sum of responses to the two cues. **(f)** Calcium responses of LPLC1 (n = 6 flies) and LPLC2 (n = 6 flies) neurons to the joint presentation of isoluminant expansion and a gray-to-white luminance change at a time offset of Δ = −250 ms, compared to the expected linear sum of responses to the two cues. **(g)** As in (e) but with an offset of Δ = 250 ms. **(h)** As in (f) but with an offset of Δ = 250 ms. **(i)** Gain in peak responses of LPLC1 and LPLC2 neurons compared to the expected linear sum as a function of temporal offsets between the two cues for gray-to-black luminance changes (gain(%) = (peak_joint_ − peak_sum_)/peak_sum_ × 100) . **(j)** As in (i) but for gray-to-white luminance changes. **(k)** Turning responses of wild-type flies (n = 21 flies) to joint presentation of isoluminant expansion and a gray-to-black luminance change at two temporal offsets (Δ = ±250 ms), as well as isoluminant expansion alone. Stimuli were presented at 90° azimuth relative to the fly’s heading. **(l)** Normalized walking speed of flies to the stimuli in (k) presented in front of the flies (n = 21 flies). Gray dots are individual flies. Vertical dotted lines mark the onset of expansion. Shaded rectangular areas represent the averaging window of 1 s after the end of expansion. Shading around curves and solid error bars represent the standard error of the mean (SEM). Stars show significance level; p-values were calculated using a paired *t*-test for (e–j) and Student’s *t*-test for (k, l) (**** p < 0.0001, *** p < 0.001, ** p < 0.01, * p < 0.05, ns p > 0.05).

Both LPLC1 and LPLC2 neurons depolarized in response to the isoluminant expanding annulus, to luminance increments, and to luminance decrements (Figures 6a to 6c). Both neurons showed greater sensitivity to luminance decrements than to luminance increments, consistent with previous reports (Klapoetke et al., 2017, 2022; Tanaka and Clark, 2022) (Figure S5). We next reasoned that if these neurons integrate these cues meaningfully during approach, the response to joint presentation of these cues should differ from the sum of their isolated responses through non-linear interactions, rather than simply reflecting their linear summation.

To test this, we measured the responses of LPLC1 and LPLC2 neurons to the joint presentation of expansion and luminance cues (Figure 6d). We presented the luminance change cue (gray-to-black or -white dot) at different temporal offsets relative to the onset of isoluminant expansion, ranging from 500 ms before to 500 ms after expansion onset (Supplemental Videos S4–S7). Both LPLC1 and LPLC2 neurons responded to this joint presentation. However, their responses depended strongly on the relative timing of the two cues. When luminance change (gray-to-black) preceded expansion onset, both neurons showed synergistic integration, responding more strongly than predicted by the linear sum of the individual cues with same temporal offsets (Figure 6e). Importantly, this supralinear integration was conserved in both neurons for luminance change from gray-to-white as well (Figure 6f). In contrast, when luminance changes followed the expansion onset, the responses to the joint presentation were sublinear or equal to the linear sum of the two cues in isolation (Figure 6g, Figure 6h). These results were consistent across a range of temporal offsets (−500 ms to +500 ms) we tested (Figure 6i, Figure 6j, Figure S6a, Figure S6b) and concord with the theory of sequential use of luminance and expansion cues (Figure 5). The presence of both supralinear and sublinear summation (Figure 6i, Figure 6j) cannot be easily explained by an indicator nonlinearity and strongly supports stimulus-dependent neural amplification and suppression. Thus, luminance changes preceding expansion yield an amplified response to expansion signals, increasing selectivity of these neurons for naturalistic approach signals. Conversely, luminance changes following expansion add little to the neural response or even suppress it (Figure 6g, Figure 6i). These findings identify luminance change as an early, complementary cue to expansion for robust approach detection. Our results from luminance-grid stimuli experiments and this temporally structured integration of luminance and expansion by LPLC1 and 2 neurons strongly support classifying them as general approach detectors in the fly brain rather than solely expansion-selective neurons.

We wanted to test whether the sequenced amplification we observed in LPLC1 and 2 was also apparent in behavior. Behavioral responses to complex visual stimuli are often the sum of responses to many features of the stimulus (Cowley et al., 2024) but we reasoned that the amplification of sequenced luminance and expansion cues might nonetheless be observable. We therefore recorded behavioral responses to the isoluminant expanding annulus and its combination with luminance changes at temporal offsets of ±250 ms. Consistent with the imaging results, flies showed a significantly stronger turning-away and slowing behaviors when the luminance change preceded the annulus expansion compared to the expansion-only control and the case when luminance change followed the expansion (Figure 6k, Figure 6l, Figure S6c, Figure S6d). Overall, these results show that the fly visual system is tuned to the natural temporal sequence of luminance and expansion cues, enhancing both the neural and behavioral response to approach.

## DISCUSSION

Expanding retinal images are canonical cues for signaling approach, yet we demonstrate that both flies and humans reliably perceive approach and retreat in purely luminance-based stimuli devoid of expansion. Luminance therefore serves not only as a complementary cue to expansion but also as a sufficient cue to infer approach in the absence of expansion. We identify LPLC1 and 2 neurons as loci where luminance and expansion cues are synergistically integrated in their natural order to generate a robust approach signal. Together, these results demonstrate how the visual system exploits the predictable dynamic structure of natural events, rather than any single cue, to achieve reliable detection.

### LPLC1 and LPLC2 integrate multiple approach cues in the fly visual system

Approaching objects present the visual system with a rich ensemble of correlated cues, including angular size, angular velocity, occlusion, contrast, and luminance change. Studies across animals have identified circuits sensitive to subsets of these features (Ache et al., 2019; Liu et al., 2011; Rind and Santer, 2004; Sun and Frost, 1998; Yilmaz and Meister, 2013). Within the fly visual system, distinct neuronal populations encode different features of object expansion: LC4 neurons are sensitive to angular speed, whereas LPLC2 neurons are sensitive to object size (Ache et al., 2019; Reyn et al., 2017). Our results demonstrate that LPLC1 and LPLC2 neurons also respond to purely luminance-based approach stimuli and serve as sites where visual cues are combined into a unified, behaviorally relevant approach signal. This parallels measurements in primate visual cortex where neurons jointly encode correlated cues, potentially facilitating perceptual inference (Bradley et al., 1995; Samonds et al., 2012). Together, these findings suggest that feature-selective neurons in the fly visual system can encode multiple correlated signatures of physical events. Their sensitivity may span a broader spectrum of temporally and spatially associated features than has yet been characterized.

### Integrating directional and non-directional inputs for approach detection

LPLC1 and LPLC2 neurons perform a sequenced integration of a non-directional luminance cue with a directional expansion cue (Figure 6), receiving direction-selective inputs from T4 and T5 cells (Klapoetke et al., 2022; Tanaka and Clark, 2022). Studies across animals, including flies, suggest that direction-selective inputs are not a general requirement for loom-sensitivity, as loom-sensitive neurons vary widely in whether they receive directional or non-directional inputs (Dunn et al., 2016; Förster et al., 2020; Jones and Gabbiani, 2010; Kim et al., 2020; Klapoetke et al., 2022; Münch et al., 2009; Rind, 1987). In primates, neurons in cortical area MST receive directional inputs and are sensitive to expanding optic flow (Tanaka et al., 1989), while LGMD neurons in locusts detect approach through a luminance-based temporal mechanism requiring no directional inputs (Fotowat and Gabbiani, 2011; Jones and Gabbiani, 2010). In larval zebrafish, loom-sensitive populations have been identified in the tectum (Dunn et al., 2016; Fotowat and Engert, 2023), while in mice looming sensitive neurons have been identified in retina and superior colliculus (Kim et al., 2020; Münch et al., 2009; Wang et al., 2021). In these systems, the specific inputs to loom-sensitive populations remain to be fully characterized. Furthermore, the nature of a neuron’s inputs does not cleanly predict its feature tuning: in flies, LPLC2 neurons receive directional inputs yet are closely tuned to object size, while LC4 neurons more closely encode angular velocity despite lacking direct directional inputs (Ache et al., 2019; Reyn et al., 2017). It is also noteworthy that neurons with non-directional inputs need not be non-directional in their computations, since non-directional inputs may be spatiotemporally delayed and amplified to give rise to directional signals (Adelson and Bergen, 1985; Borst and Egelhaaf, 1989). Together, these observations suggest that the relationship between input directionality, feature tuning, and loom detection strategy remains complex. Our results provide a concrete example of how directional and non-directional cues can be synergistically combined in a single circuit.

### Sequenced integration of luminance and expansion cues

The sequenced cue integration in LPLC1 and LPLC2 reflects a broader motif in neural computations that exploit natural temporal cue structures. In audiovisual integration, visual input has been reported to reshape auditory cortex responses when it precedes the audio input by 20–80 ms (Kayser et al., 2008). This temporal window mirrors the natural precedence of visual over auditory signals in the physical world, where light travels faster than sound across behaviorally relevant distances. Direction selectivity in motion-processing circuits provides a canonical example where spatiotemporally offset inputs in neighboring locations amplify or suppress responses to preferred and null direction motion (Adelson and Bergen, 1985; Barlow and Levick, 1965; Hassenstein and Reichardt, 1956). Here, we demonstrate that luminance change preceding expansion enhances the responses of approach-sensitive neurons. These results are consistent with a general principle in which specific sequences of stimuli are enhanced via gain changes, allowing neural circuits to exploit the stereotyped temporal structure of natural sensory events.

### Contrast polarity of approach signals

Many studies across animals have found that dark looming stimuli elicit stronger behavioral and physiological responses than light looming stimuli (Dewell et al., 2022; Kim et al., 2020; Klapoetke et al., 2017, 2022; Lee et al., 2020; Rind and Simmons, 1992; Temizer et al., 2015; Vries and Clandinin, 2012; Yilmaz and Meister, 2013). It is therefore notable that, while LPLC1 and LPLC2 neurons show the same asymmetry to luminance increments and decrements (Figures S5, 6a and 6b) and to dark and light looming discs (Klapoetke et al., 2022), responses to expansion in both neurons were enhanced by both luminance increments and decrements (Figures 6e and 6f). Ecologically, this is unsurprising: approaching objects could be either lighter or darker than the background, and an approach detector should be sensitive to either. Indeed, humans perceive objects as nearer when they have higher contrast against the background, regardless of polarity (Dosher et al., 1986; Schwartz and Sperling, 1983). The overall amplitude response differences to light vs. dark approaching objects may still represent a tuning to the natural statistics of approach.

While luminance increments and decrements both enhance responses to isoluminant expansion, the luminance-grid illusory approach and retreat percepts clearly rely on luminance asymmetry (Figures 1b, 1d and 2k). The illusory percept requires the grid (Weiss et al., 2004) and is thus likely the result of complex spatiotemporal integration. Because the two sawtooth waveforms (ramp-up and ramp-down) carry luminance derivatives of equal magnitude but opposite sign, which average to zero, linear derivative-based detection mechanisms would fail to distinguish between them. The luminance-grid illusion therefore seems to arise from an imbalance in the processing of contrast increments and decrements, even though both polarities enhance approach detection when combined with expansion (Figure 6). Similar imbalances may underlie other illusions, including the peripheral drift illusion (Fraser and Wilcox, 1979), where purely static spatial patterns elicit directional motion percepts due to asymmetric ON/OFF processing (Agrochao et al., 2020). In motion detection, light–dark asymmetries are also apparent in humans and flies (Chen et al., 2019; Clark et al., 2014; Hu and Victor, 2010; Leonhardt et al., 2016; Wu et al., 2025), where the asymmetry is proposed to arise as a tuning to light–dark asymmetries in natural scenes (Fitzgerald and Clark, 2015; Fitzgerald et al., 2011).

### Parallel percepts of luminance-based approach and retreat in flies and humans

Flies and humans appear to share the same percepts of approach and retreat in the luminance-grid stimuli (Figures 1 to 3). This parallel strongly suggests that core computational principles for approach and retreat detection are shared across phylogenetically distant species. Despite radically different neural architectures, both visual systems extract approach signals from temporally asymmetric luminance modulation and exhibit weakened percepts under symmetric luminance modulation in the grid. These parallel computations suggest there are fundamental constraints on how visual systems infer three-dimensional approach from two-dimensional, retinal inputs. While the natural statistics of approach are not well understood, we propose that the luminance-grid illusion ultimately arises from light-dark imbalances in processing of naturalistic approach statistics, which also include light-dark asymmetries of natural scenes.

### The temporal ordering of approach cues is a geometric inevitability

Physical principles constrain the dynamics of approaching objects, and therefore the geometry of approach cues is conserved across sighted animals. The temporal order of luminance and expansion is also constrained in a broad sense: expansion only becomes informative late in approach because spatial resolution and geometry both limit expansion signals at long distances, whereas luminance and contrast signals may remain informative at all times. The temporal order of these cues is therefore a geometric inevitability rather than a species-specific adaptation. Because of this, the mechanism of sequenced integration of luminance and expansion we have identified may apply broadly to approach sensitive circuits in other species to generate reliable approach signals. Luminance and expansion are ultimately experimentally-derived analytical distinctions: a real approaching object produces many cues as a single, unified, temporally structured event. The sensitivity of approach sensors to these features can be viewed as tuning these neurons to the natural spatiotemporal signature of approach itself rather than interactions between independent features.

More broadly, the findings here highlight how the temporal structure of events in the world organizes circuit computations by shaping cue integration strategies for reliable perception.

## ACKNOWLEDGMENTS

We thank members of the Clark lab for useful input and discussion. We also thank Sam McDougle for help with human psychophysics and Brian Scholl for helpful discussions. NM was supported by CAPES fellowship. DAC and this project were supported by NIH R01 EY026555.

## AUTHOR CONTRIBUTIONS

Experiments were conceived by HV, NM, and DAC. HV and NM collected behavioral data. HV and HW collected imaging data. HV, NM, and HW interpreted and analyzed data. The manuscript was drafted by HV and edited by HV and DAC.

## GENERATIVE ARTIFICIAL INTELLIGENCE STATEMENT

During the preparation of this work the authors used Claude Sonnet 4.6 (Anthropic, USA) in order to occasionally improve grammar, readability, and word choice. After using this tool, the authors reviewed and edited the content as needed and take full responsibility for the content of the publication.

## MATERIALS AND METHODS

### Data and code availability

All data and code to create the figures in this paper will be made available on public repositories upon publication.

### Human psychophysics experiments

Human psychophysics experiments were conducted following protocols approved by the Institutional Review Board of Yale University and in accordance with the Declaration of Helsinki. Participants provided written informed consent prior to testing and were compensated for their participation. All stimuli for human psychophysics experiments were generated using MATLAB (MathWorks, Natick, MA, USA) and Psychtoolbox (Brainard and Brainard, 1997; Kleiner et al., 2007; Pelli, 1997) and presented on a monitor (refresh rate: 60 Hz and screen gamma of 1.8) at a viewing distance of 57 cm in a dimly lit, sound-attenuated room. Luminance-grid stimuli and their different variations were constructed by arranging a 6×6 grid of circles with diameter 2.5°. Initial circle luminances were chosen between 0 and 1 from a uniform distribution and then modulated either with pure sawtooth waves or their interpolations. All stimulus waveforms had a period of 0.5 s. Ramp-up and ramp-down stimuli, or their interpolated versions, were presented for a duration of 2 s in randomized order, interleaved by a 2 s judgment window. Before the beginning of the experiments, participants were verbally instructed to judge whether the stimulus appeared to be approaching or retreating, with instructions similar to: “You will see a series of stimuli. After each presentation, please indicate whether the stimulus appeared to approach or retreat. Use the keys provided to give your response.” Participants indicated their responses via button press, with reaction times and choices recorded on each trial.

### Fly husbandry

Flies were reared at ∼50% relative humidity on a glucose-based food (Archon Scientific, Durham, USA; recipe D20102). Flies were reared at 20°C for behavior experiments and 25°C for imaging experiments in growth incubators with fixed 12-hr:12-hr light:dark circadian cycles. For all behavior and imaging experiments, non-virgin female flies were collected using CO_2_ anesthetization within 18–24 h of eclosion and then used 12 h later for the experiments. All behavioral experiments were performed within 3-h windows after lights-on or before lights-off. The lines used in the experiments are summarized in the tables below.

#### Reagent or Resource

| Experimental Models: Organisms/Strains | Source | Identifier |
| --- | --- | --- |
| *D. melanogaster*: +; +; + (IsoD1 wild type) | (Gohl et al., 2011) | N/A |
| *D. melanogaster*: +; UAS-shibire; UAS-shibire (shibire) | (Kitamoto, 2001) | N/A |
| *D. melanogaster*: +; UAS-GCaMP6f; + (GCaMP6f) | (Chen et al., 2013) | N/A |
| *D. melanogaster*: w; R64G09-AD; R37H04-DBD (LPLC1) | (Wu et al., 2016) | JRC: #OL0029B |
| *D. melanogaster*: w; R19G02-AD; R75G12-DBD (LPLC2) | (Wu et al., 2016) | JRC: #OL0048B |
| *D. melanogaster*: w; R19G02-AD; R12E04 (LPLC2) | (Klapoetke et al., 2017) | JRC: #OL0047B |
| *D. melanogaster*: w; R22H02-AD; R20G06-DBD (LC11) | (Wu et al., 2016) | BDSC: #68362 |
| *D. melanogaster*: w; empty-AD; empty-DBD (empty split Gal4) | (Hampel et al., 2015) | BDSC: #79603 |

#### Genotypes Used

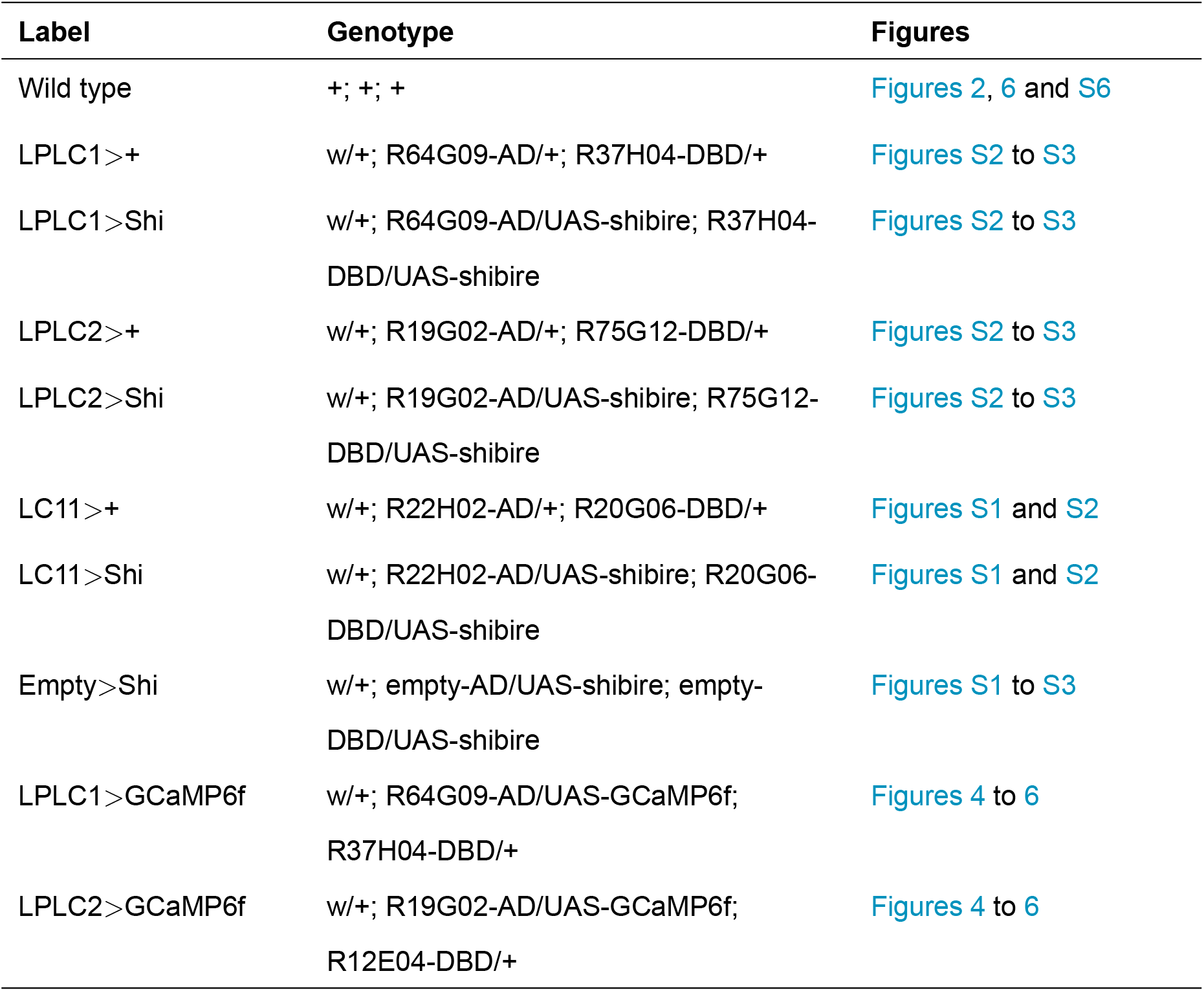

### Stimulus

All visual stimuli used in this study were generated using MATLAB (MathWorks, Natick, MA, USA) and Psychtoolbox (Brainard and Brainard, 1997; Kleiner et al., 2007; Pelli, 1997). Stimuli were displayed on a virtual cylinder surrounding the fly using a LightCrafter DLP (Texas Instruments, Dallas, Texas, USA) operating at a refresh rate of 180 frames per second. The stimuli were projected using green light (peak 520 nm; mean intensity ∼100 cd/m^2^). The screen geometry was as described previously (Creamer et al., 2019).

#### Stimuli

Stimulus Stimuli
| Stimulus | Description | Figures |
| --- | --- | --- |
| Ramp-up and ramp-down luminance-grid stimuli for human psychophysics | Luminance-grid stimuli consisting of a 6×6 grid of circles with diameter 2.5°. Initial circle luminances were chosen between 0 and 1 from a uniform distribution and then modulated with sawtooth waves of period 0.5 s. | Figures 1 and S1 |
| Ramp-up and ramp-down luminance-grid stimuli for fly psychophysics | Luminance-grid stimuli consisting of a 6×6 grid of circles with diameter 10°. Initial circle luminances were chosen between 0 and 1 from a uniform distribution and then modulated with sawtooth waves of period 0.5 s. | Figures 1 to S3 |
| Expanding disc stimuli | Circular discs (initial radius 5°) were presented at 90° and 0° azimuth relative to the fly's heading on a gray background. After 2 s, the discs expanded to a final radius of 35° and then remained static for 1 s before disappearing. | Figures S1 to 3 |
| Contracting disc stimuli | Circular discs (initial radius 35°) were presented at 90° and 0° azimuth relative to the fly's heading on a gray background. After 2 s, the discs contracted to a final radius of 5° and then remained static for 1 s before disappearing. | Figures S1 to S3 |
| Expanding and contracting disc stimuli for imaging | Similar to behavior, but the discs expanded from an initial radius of 0° to a final radius of 30° and then remained static for 1 s. | Figures 4 and S4 |
| Isoluminant expansion stimulus | An isoluminant annulus with an initial inner radius of 5°, an outer radius of 15°, and a checkerboard pattern with check size 5° × 22.5° (radial visual angle × annulus angular coordinate) was presented at the receptive-field center. The inner and outer radii expanded at 30°/s until the outer radius reached 30°. | Figure 6a |
| Luminance change stimulus | A circular dot (radius 2°, 5°, 10°, 15°, 20°) was presented at the receptive-field center and changed luminance from gray to white or from gray to black. | Figure S5 |
| Joint-cue stimulus | The luminance change was presented using a circular dot (radius 5°) at the receptive-field center at different time offsets relative to isoluminant annulus expansion onset. | Figures 6e to 6j |
| Joint-cue stimulus for behavior | As in imaging, but the isoluminant annulus expanded from an inner radius of 5° to 35° at 60°/s. The stimulus was presented either in front of the fly or at 90° azimuth relative to the fly's heading. | Figures S6, 6k and 6l |

### Fly behavioral experiments

Behavioral experiments were performed following the procedures established in previous studies (Creamer et al., 2019; Tanaka and Clark, 2022). Briefly, flies were cold-anesthetized and then tethered to surgical needles with UV-curable epoxy on their dorsal thorax before being positioned atop air-supported balls. Visual stimuli were projected onto a surrounding panoramic screen covering 270° azimuth and 106° elevation and the fly walking responses were quantified by measuring the rotation of the ball. The rig temperature was maintained at ∼34°C to promote locomotion and activate thermogenetic tools.

#### Behavior data analysis

Walking speed was normalized to the mean speed during the 500 ms pre-stimulus window. Normalized speed and turning angular-velocity traces were averaged across stimulus presentations. Mirror-symmetric stimulus pairs were combined for walking and turning traces. Individual mean traces were temporally averaged for statistical analysis over the durations noted in figure captions. Frozen fraction was calculated by counting the number of flies whose normalized walking speed dropped to less than or equal to zero and dividing by the total number of flies. Group means and standard errors were derived from individual mean traces to characterize response dynamics.

### Two-photon calcium imaging

Two-photon calcium imaging was performed following protocols described previously (Tanaka and Clark, 2022). One-to two-day-old female flies were cold-anesthetized and head-fixed to a custom-made round metal holder using UV-cured epoxy. The back of the head was exposed by cutting the cuticle and removing fat tissue over the optic lobe. The holder was filled with oxygenated sugar-saline solution before transfer to the microscope. Neural imaging was performed using a two-photon microscope (Scientifica, Clarksburg, NJ, USA) with a 20× water-immersion objective (Olympus, Waltham, MA, USA). Visual stimuli were projected onto a panoramic screen covering 270° azimuth and 69° elevation of the fly’s visual field using a DLP projector (Texas Instruments, Dallas, Texas, USA). To account for head tilt in the holder, all visual stimuli were pitched 45° forward relative to the screen. Green-light bleed from the projector into the PMT was prevented using a 565/24 filter in series with a 560/25 filter on the projector (Semrock, Rochester, NY, USA). PMT input was filtered using a 512/25 filter to capture green fluorophore emissions. A femtosecond Ti:sapphire laser (Spectra-Physics, San Jose, CA, USA) was used to excite the sample at 930 nm. Laser power at the sample was maintained below 40 mW. All imaging data were acquired at ∼8.5 Hz using ScanImage software (Pologruto et al., 2003). A probe was presented at the start and end of each recording to identify responsive ROIs and verify fly viability at the end of the experiment.

### Calculation of calcium responses

ROIs were defined manually or via a watershed segmentation using time-averaged fluorescent images. Stimulus bleed-through was removed from each frame by subtracting a pixel-averaged signal computed in a background region, identified as the largest contiguous region in the time-averaged frame below the 10th percentile in brightness. Fluorescent traces were converted to Δ*F* /*F* to control for expression variability and fluorophore photobleaching. Baseline fluorescence (*F*) was obtained by averaging fluorescence across pixels within each ROI, then fitting a decaying exponential *A* exp(−*t*/*τ*) to time-averaged fluorescence during interleave epochs across each recording of 5–10 min, with *τ* constrained to be uniform across all ROIs in each recording. This baseline fluorescence (*F*) was subtracted from raw ROI fluorescence traces, and the residual (Δ*F*) was divided by the baseline fluorescence (*F*) to yield Δ*F* /*F* traces. Responses to identical stimulus repetitions were averaged within ROIs, then across all ROIs per fly to generate individual mean responses for each fly. Small fluctuations in Δ*F* /*F* over time were removed by subtracting time-averaged Δ*F* /*F* during the 500 ms pre-stimulus window. For statistical comparisons across conditions and genotypes, mean or peak responses were calculated over appropriate windows noted in the figure captions.

### Receptive field mapping

For single-cell recordings (Figures 4 to 6), each stalk’s receptive field (RF) was mapped prior to stimulus presentation following methods described previously (Tanaka and Clark, 2022), with subsequent stimuli centered on the estimated RF location. Initial RF approximation was determined interactively using small translating black squares. A 10° black square then swept a 40° × 40° region surrounding the approximate location at 60°/s, moving horizontally and vertically in 5° increments. Neural responses (Δ*F* /*F*) were averaged temporally over a 1.5 s window from stimulus onset and across motion directions for each azimuth and elevation, yielding horizontal and vertical spatial tuning curves. Independent Gaussian fits to these curves provided mean values that served as the estimated RF center. Only RFs with Gaussian fits exceeding *R*^2^ > 0.8 were included in the analyses.

### Motion detection simulation

We simulated responses of a simplified motion-detecting pathway (T4/T5-like) to a one-dimensional looming-edge stimulus using a previously published minimal synaptic model of T4 response properties (Zavatone-Veth et al., 2020).

The visual field was represented as a one-dimensional angular axis spanning 0–180° with spatial resolution of 0.05°. The looming stimulus was a binary box-car function (“object”) centered at 90°. For each simulation, looming dynamics were parameterized by the size-to-speed ratio (*R*/*v*), where *R* is the half-size of the object and *v* is its approach speed. The apparent object half-angle followed:

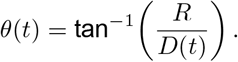

where *D* is the relative distance between the detector and the object.

Following prior modeling of the motion detector responses (Zavatone-Veth et al., 2020), to mimic optical blur, the stimulus was convolved in space with a Gaussian kernel of full-width-at-half-maximum 5.7°. Temporal filtering was applied independently at each spatial position using a low-pass filter (*f*) with time constant *τ* = 0.10 s and a high-pass filter equal to the derivative form of *f* with *τ* = 0.10 s.

The T4 architecture was implemented by spatially shifting the low-pass output by a photoreceptor spacing of 5° to create “leading” and “trailing” channels. Each photoreceptor channel was half-wave rectified and weighted to produce conductances. Two mirror-symmetric subunits were computed and tiled across the one-dimensional space. A calcium proxy was then generated by half-squaring (squaring after half-wave rectification) and summing the two mirror-symmetric subunits.

To obtain an estimate of the integrated population response, *Q*(*t*), for a two-dimensional disc, we summed the one-dimensional radial calcium proxy across space (*x* ∈ [0, *θ*(*t*)]) and scaled it by a factor of 2*πθ*(*t*) to account for radial signals around the circumference of the disc. The reference detector response was computed at a fixed object diameter of 40°, chosen to mimic the typical receptive field size of loom sensors (Klapoetke et al., 2022; Tanaka and Clark, 2022), evaluated across all *R*/*v* values. A scalar detection threshold (Θ) was then set as a fraction of this maximum reference response. For each *R*/*v*, the detection time was defined as the time index at which *Q*(*t*) ≥ Θ.

## APPENDIX 1

## Rate of change of object size during approach

Consider a circular disc of radius *R* approaching the observer along the line of sight with constant speed *v* > 0. Let *D*(*t*) denote the distance from the observer to the disc center, so *D*(*t*) = *D*_0_ − *vt*. The half-angle subtended by the disc is

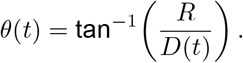

For 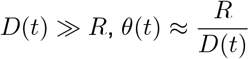 and the rate of change of apparent size is

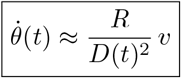

### Irradiance at the retina due to object luminance

Consider a small receiver (retina point) at axial distance *D* from a Lambertian disc of radius *R* and uniform luminance *L*_*d*_, presented on a uniform background of luminance *L*_0_. Let Ω_*d*_ denote the solid angle subtended by the disc at the receiver. The incident radiance is

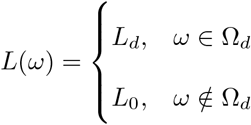

The hemispherical irradiance at the receiver is (Lambert cosine law)

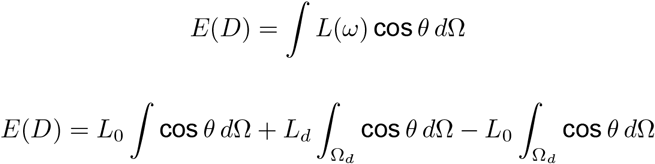

Let Δ*L* ≡ *L*_*d*_ − *L*_0_,

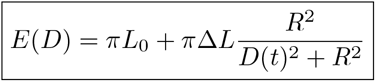

For 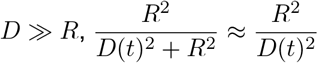,

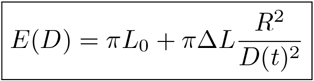

If the disc approaches at constant speed *v* > 0 so that 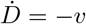, then

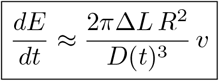

The rate of change of apparent size of an approaching object is proportional to distance as 1/*D*(*t*)^2^, while the rate of change of irradiance at the detector due to the object’s luminance is proportional to distance as 1/*D*(*t*)^3^. This means that as an object approaches, the irradiance signal grows faster than the apparent size signal and could be detected before any detectable expansion occurs.

### Total motion signals from an approaching disc

For motion signals proportional to the velocity, the net motion signal grows with both the length of the moving edge (*C*) and the rate of change of edge. The length of moving edge is proportional to the apparent circumference of the approaching disc:

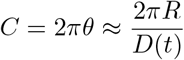

while the change rate for each edge is

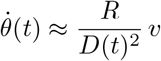

therefore, the net motion signal is proportional to:

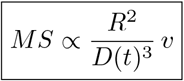

This means that as an object approaches, the irradiance signal and the total motion signal grow at the same rate with respect to *D*. However, the motion signal requires the object to subtend a minimum angular size before sufficient edge units are recruited for detection. In contrast, the irradiance signal may be detectable even when the object occupies a very small visual angle as long as Δ*L* is sufficiently large. This suggests that irradiance could serve as an early warning signal, prior to the object reaching the angular size necessary to drive a motion response.

---

**Figure S1:**
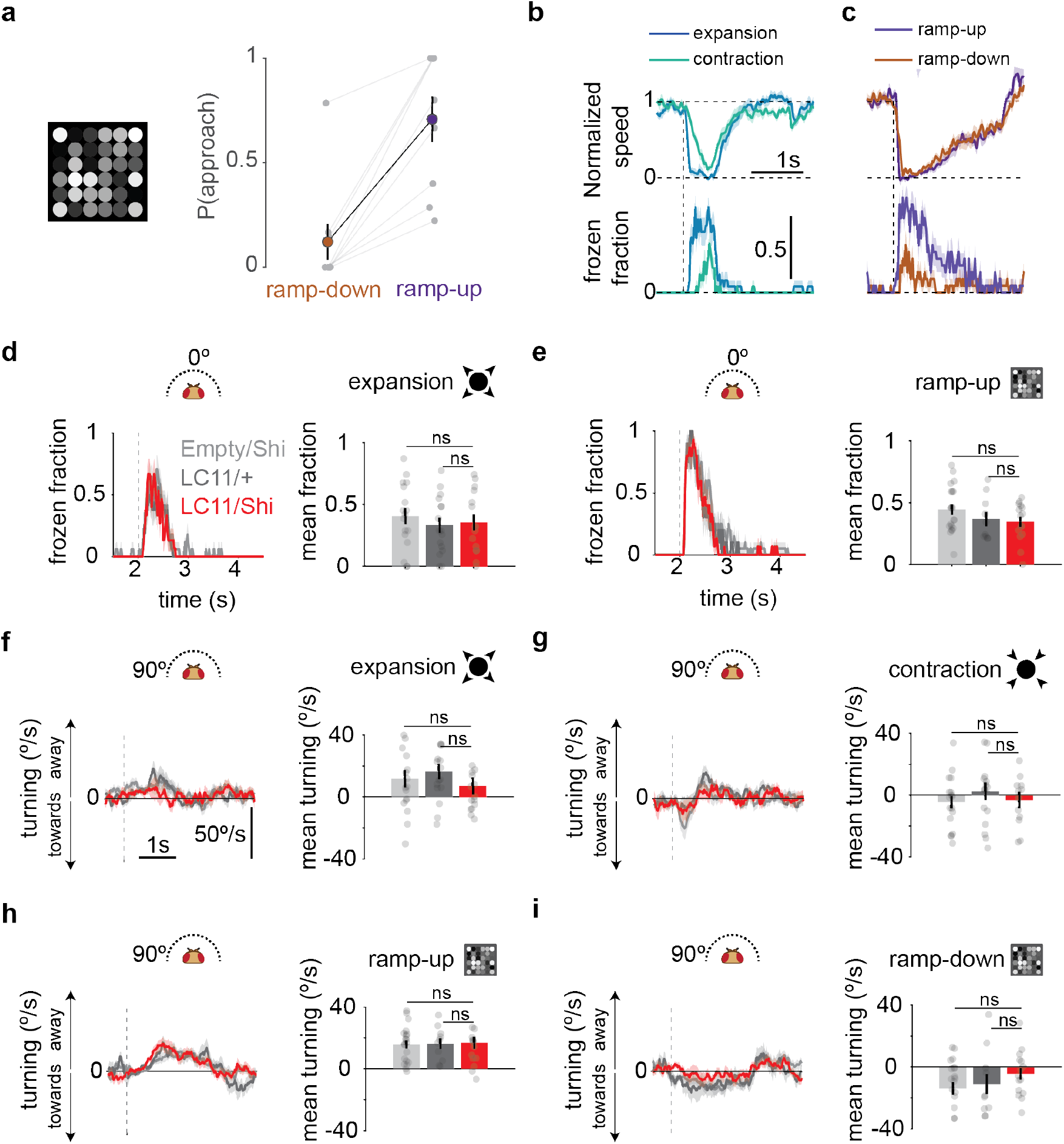
Human psychophysics additional data and LC11 silencing effects on fly behavior. **(a)** Probability of approach perception in human subjects (n = 9) for luminance-grid stimuli presented on a gamma-corrected monitor. Linked gray dots are individual subjects. **(b)** Normalized walking speed and corresponding frozen fraction in response to expanding/contracting disc stimuli (60°/s) (n = 20 flies). **(c)** As in (b) but for ramp-up and ramp-down stimuli (period = 0.5 s) (n = 17 flies). **(d)** Silencing LC11 neurons has no effect on the fly’s freezing response to an expanding disc stimulus. Empty/Shi (n = 17 flies), LC11/+ (n = 16 flies), LC11/Shi (n = 15 flies). **(e)** Silencing LC11 neurons has no effect on the fly’s freezing response to a ramp-up stimulus. Empty/Shi (n = 20 flies), LC11/+ (n = 10 flies), LC11/Shi (n = 15 flies). Mean frozen fraction was calculated in a 1 s window after stimulus onset for both stimuli. **(f)** Silencing LC11 neurons has no effect on the fly’s turning response to expansion. **(g)** Silencing LC11 neurons has no effect on the fly’s turning response to contraction. **(h)** As in (f) but for ramp-up luminance-grid stimuli. **(i)** As in (h) but for ramp-down luminance-grid stimuli. Mean turning for expansion and contraction was calculated in a 1 s window after stimulus onset, and in the last 1 s for luminance-grid stimuli. Gray dots are individual flies. Vertical dotted lines mark the onset of expansion (contraction) and luminance-grid modulation. Shading around curves and error bars in the bar plots represent the standard error of the mean (SEM). Stars show significance level; p-values were calculated using Student’s *t*-test (**** p < 0.0001, *** p < 0.001, ** p < 0.01, * p < 0.05, ns p > 0.05).

**Figure S2:**
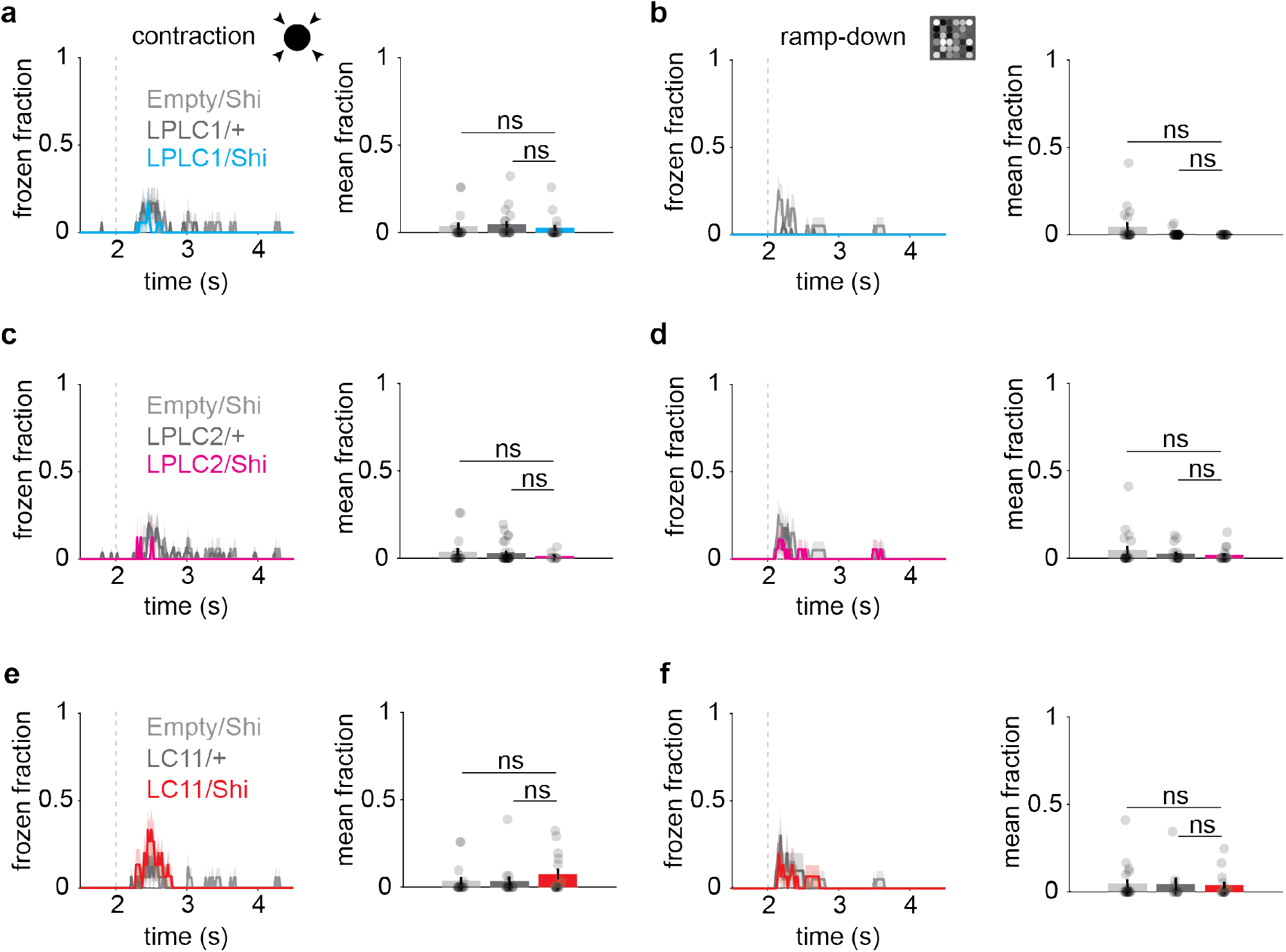
Flies show minimal freezing in response to contraction and ramp-down stimuli. **(a)** Silencing LPLC1 neurons has no effect on the fly’s freezing response to a contracting disc stimulus. Plots are as in Figure 3. Empty/Shi (n = 17 flies), LPLC1/+ (n = 18 flies), LPLC1/Shi (n = 18 flies). **(b)** Silencing LPLC1 neurons has no effect on the fly’s freezing response to a ramp-down stimulus. Empty/Shi (n = 20 flies), LPLC1/+ (n = 29 flies), LPLC1/Shi (n = 15 flies). **(c)** Silencing LPLC2 neurons has no effect on the fly’s freezing response to a contracting disc stimulus. Empty/Shi (n = 17 flies), LPLC2/+ (n = 18 flies), LPLC2/Shi (n = 8 flies). **(d)** Silencing LPLC2 neurons has no effect on the fly’s freezing response to a ramp-down stimulus. Empty/Shi (n = 20 flies), LPLC2/+ (n = 17 flies), LPLC2/Shi (n = 18 flies). **(e)** Silencing LC11 neurons has no effect on the fly’s freezing response to a contracting disc stimulus. Empty/Shi (n = 17 flies), LC11/+ (n = 16 flies), LC11/Shi (n = 15 flies). **(f)** Silencing LC11 neurons has no effect on the fly’s freezing response to a ramp-down stimulus. Empty/Shi (n = 20 flies), LC11/+ (n = 10 flies), LC11/Shi (n = 15 flies). Gray dots are individual flies. Vertical dotted lines mark the onset of contraction and luminance-grid modulation. Shading around curves and error bars in the bar plots represent the standard error of the mean (SEM). Stars show significance level; p-values were calculated using Student’s *t*-test (ns p > 0.05).

**Figure S3:**
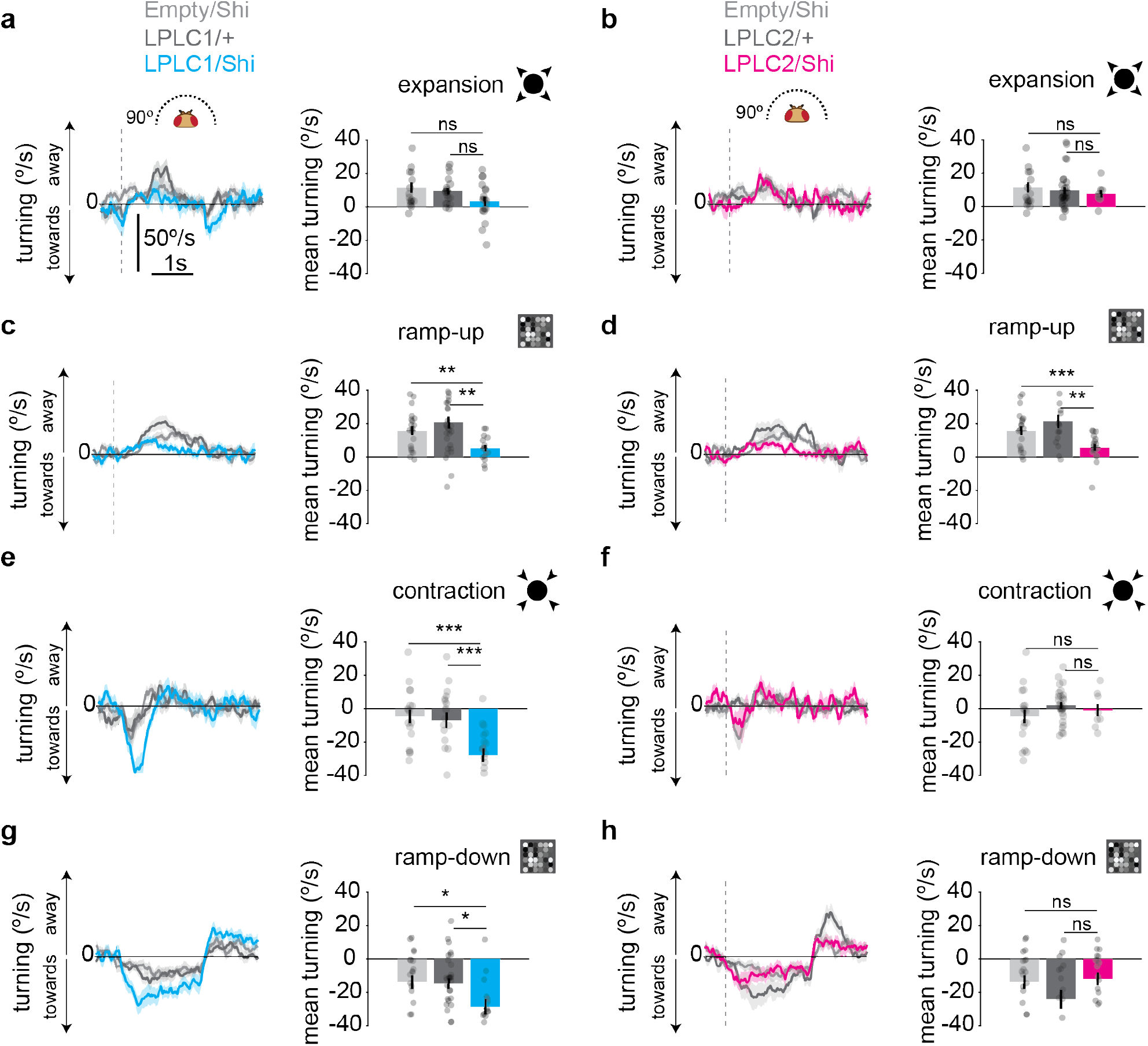
Silencing LPLC1 and LPLC2 neurons differentially affects fly turning behavior. **(a)** Turning responses to an expanding disc stimulus (60°/s) with LPLC1 neurons silenced. Empty/Shi (n = 17 flies), LPLC1/+ (n = 18 flies), LPLC1/Shi (n = 18 flies). **(b)** Turning responses to an expanding disc stimulus (60°/s) with LPLC2 neurons silenced. Empty/Shi (n = 17 flies), LPLC2/+ (n = 18 flies), LPLC2/Shi (n = 8 flies). **(c)** Turning responses to a ramp-up stimulus (period = 0.5 s) with LPLC1 neurons silenced. Silencing LPLC1 neurons significantly reduced the turning away response. Empty/Shi (n = 20 flies), LPLC1/+ (n = 29 flies), LPLC1/Shi (n = 15 flies). **(d)** Turning responses to a ramp-up stimulus (period = 0.5 s) with LPLC2 neurons silenced. Silencing LPLC2 neurons significantly reduced the turning away response. Empty/Shi (n = 20 flies), LPLC2/+ (n = 17 flies), LPLC2/Shi (n = 18 flies). **(e)** As in (a) but for a contracting disc stimulus. Silencing LPLC1 significantly increased turning toward the contracting disc stimulus. **(f)** As in (b) but for a contracting disc stimulus. **(g)** Turning responses to a ramp-down stimulus with LPLC1 neurons silenced. Silencing LPLC1 significantly increased turning toward the ramp-down stimulus. **(h)** Turning responses to a ramp-down stimulus with LPLC2 neurons silenced. Mean turning for expansion and contraction was calculated in a 1 s window after stimulus onset, and in the last 1 s for luminance-grid stimuli. Gray dots are individual flies. Vertical dotted lines mark the onset of expansion and luminance-grid modulation. Shading around curves and error bars in the bar plots represent the standard error of the mean (SEM). Stars show significance level; p-values were calculated using Student’s *t*-test (**** p < 0.0001, *** p < 0.001, ** p < 0.01, * p < 0.05, ns p > 0.05).

**Figure S4:**
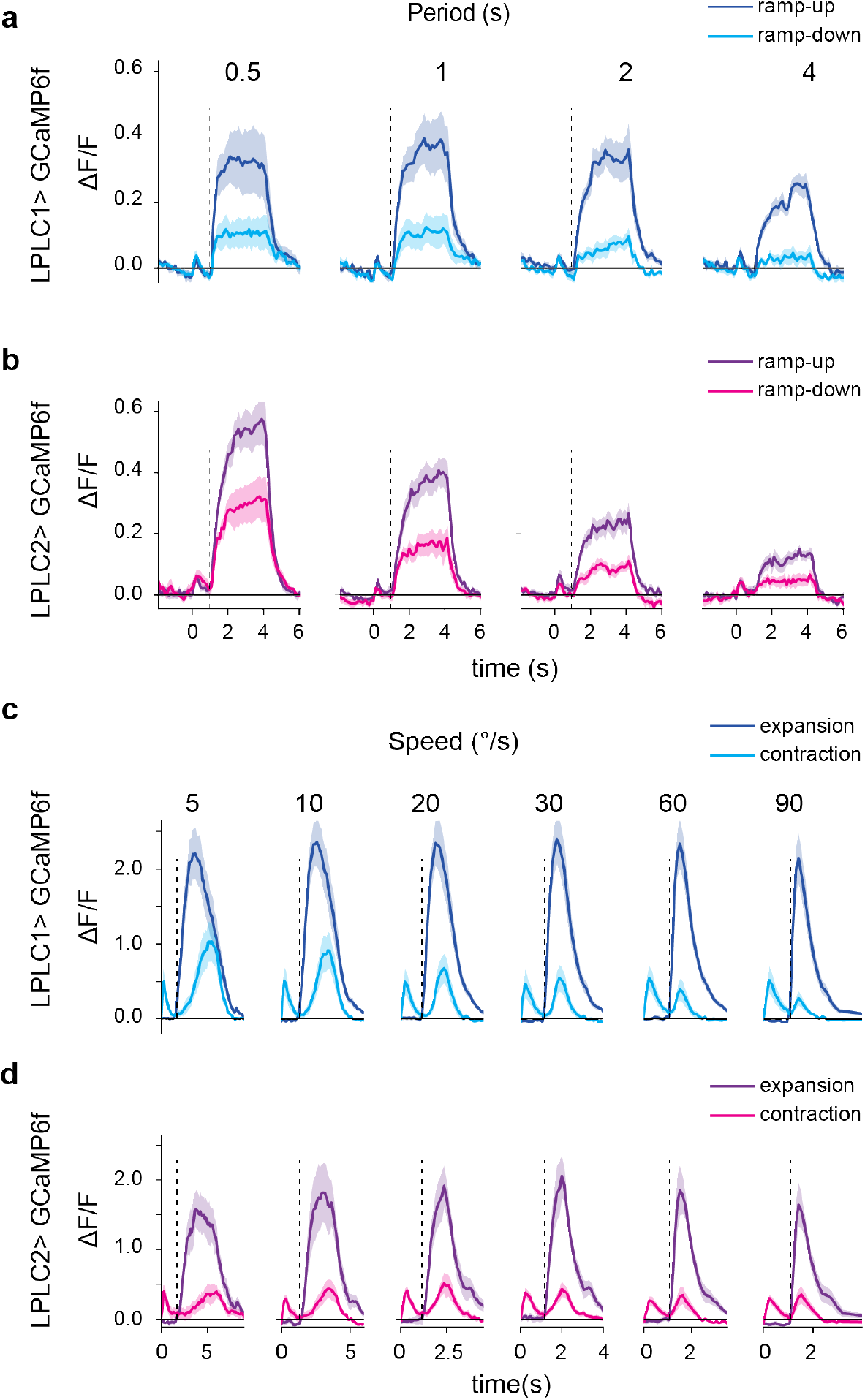
LPLC1 and LPLC2 neurons respond differentially to ramp-up and ramp-down luminance-grid stimuli and to expanding and contracting disc stimuli. **(a)** Calcium responses of LPLC1 neurons (n = 7 flies) to ramp-up and ramp-down luminance-grid stimuli presented at four sawtooth periods. **(b)** As in (a) but for LPLC2 neurons (n = 8 flies). Both neuron types exhibit high selectivity for ramp-up over ramp-down stimuli across all sawtooth periods. **(c)** Calcium responses of LPLC1 neurons (n = 8 flies) to expanding and contracting disc stimuli presented at six speeds. **(d)** As in (c) but for LPLC2 neurons (n = 6 flies). Both neuron types exhibit high selectivity for expansion over contraction stimuli across all speeds.

**Figure S5:**
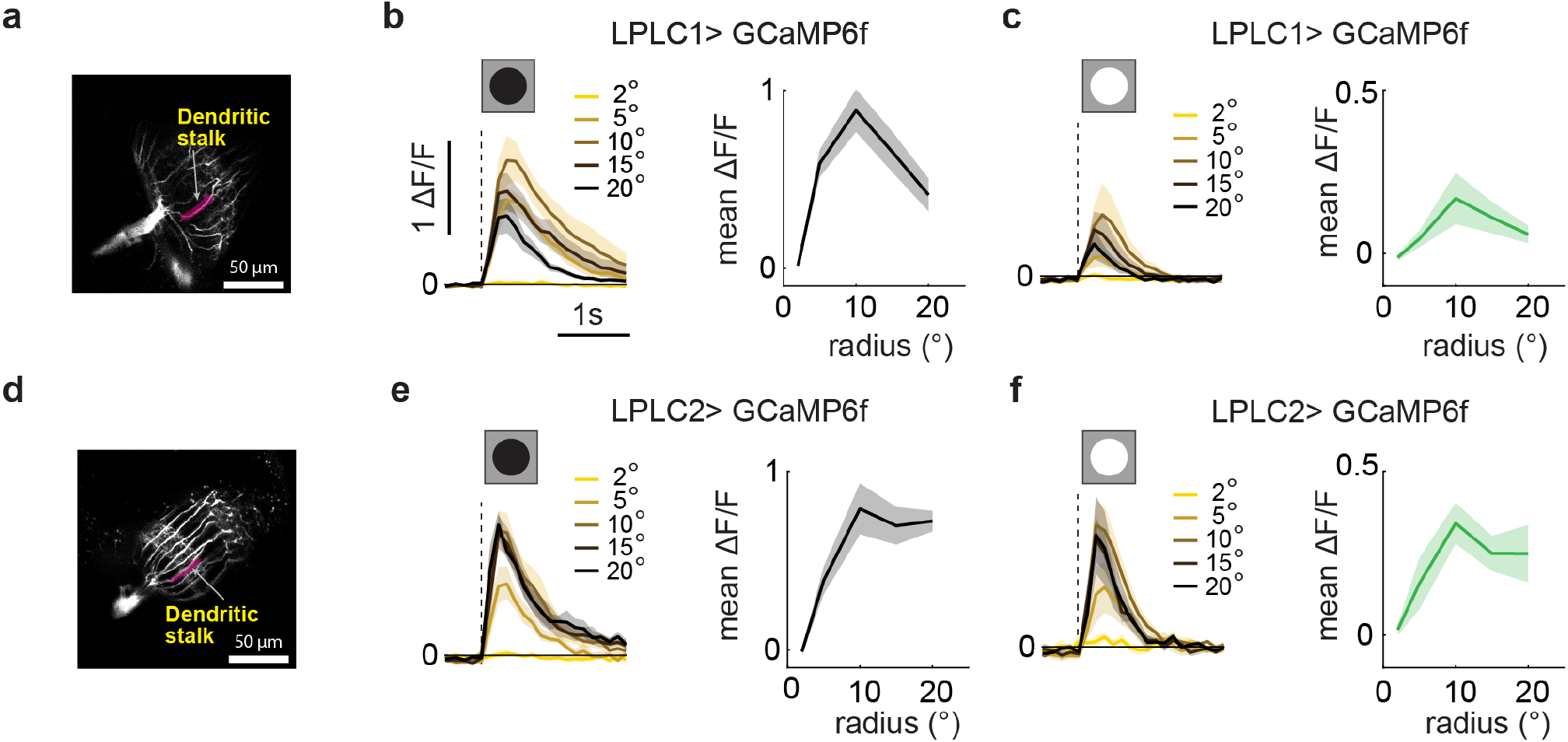
LPLC1 and LPLC2 neurons respond to gray-to-black or gray-to-white transitions in a size dependent manner. **(a)** Two-photon image of LPLC1 population expressing calcium indicator GCaMP6f. **(b)** Calcium responses of LPLC1 neurons (n = 6 flies) to gray-to-black dots of different radii presented at their receptive field (RF) centers. LPLC1 neurons depolarize in response to gray-to-black dot transition in a size dependent manner. **(c)** As in (b) but with gray-to-white dot transitions. **(d)** Two-photon image of LPLC2 population expressing calcium indicator GCaMP6f. **(e)** Calcium responses of LPLC2 neurons to gray-to-black dots of different sizes presented at their receptive field (RF) centers. LPLC2 neurons depolarize in response to gray-to-black dot transition in a size dependent manner. **(f)** Same as (e) but with gray-to-white dot transitions. Mean responses were calculated in a 2 s window after dot onset. Vertical dotted lines mark the onset of the transition. Shaded areas represent the standard error of the mean (SEM).

**Figure S6:**
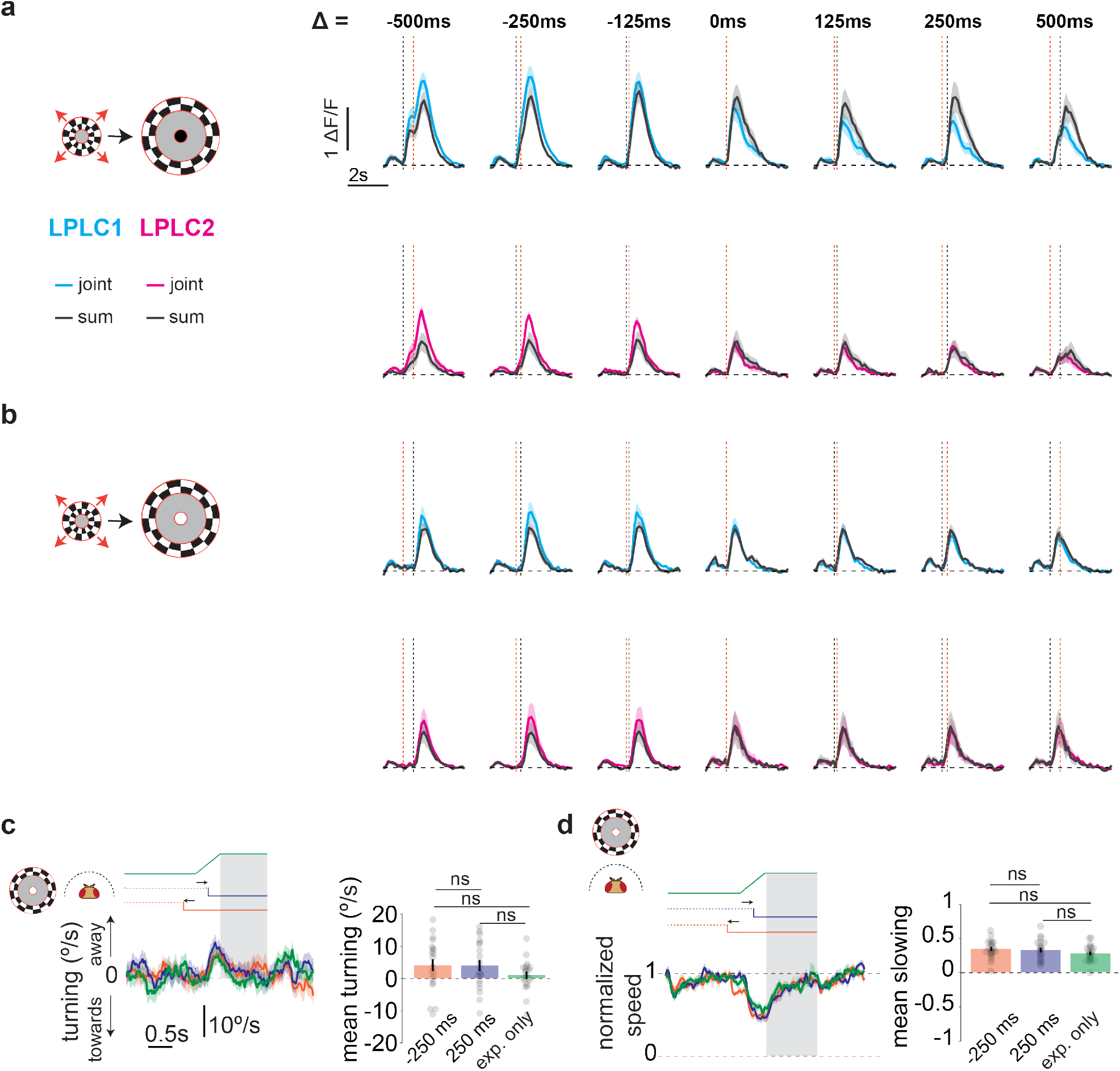
Calcium responses of LPLC1/2 neurons and fly behavior to joint cue stimuli. **(a)** Calcium responses of LPLC1 (n = 6 flies) and LPLC2 (n = 6 flies) neurons to the joint presentation of isoluminant expansion and gray-to-black luminance change at various time offsets ranging between −500 ms to +500 ms compared to the expected linear sum. **(b)** As in (a) but with gray-to-white luminance change. **(c)** Turning responses of wild-type flies (n = 20 flies) to joint presentation of isoluminant expansion and gray-to-white luminance change at two temporal offsets (Δ = ±250 ms), as well as to isoluminant expansion alone. The stimuli were presented at 90° azimuth relative to the fly’s heading. Mean turning responses in all conditions were not significantly different. **(d)** Normalized walking speed of flies in response to the cue combinations in (a) presented in front of the fly. Similar to turning, no two conditions were significantly different. Vertical dotted lines mark the onset of expansion. Shaded rectangular areas represent the averaging window of 1 s after the end of expansion. Shading around curves and error bars in the bar plots represent the standard error of the mean (SEM). p-values were calculated using a paired *t*-test (ns p > 0.05).

**Supplementary Video S1. Luminance-grid stimuli generate approach and retreat percepts in humans**. Video shows a 6×6 grid of circles whose luminance is modulated with either an increasing sawtooth (ramp-up) or decreasing sawtooth (ramp-down) waveform (period 0.5 s) for 2 s each, repeated three times (12 s total). Frame rate: 60 fps. Adapted from stimulus at URL: https://michaelbach.de/ot/sze-lumiloom/.

**Supplementary Video S2. Luminance-grid stimuli with balanced luminance increments and decrements**. Video shows a 6×6 grid of circles whose luminance is modulated with a symmetrical triangular waveform (period 0.5 s) for 4 s. Frame rate: 60 fps.

**Supplementary Video S3. Isoluminant expansion**. Video shows isoluminant annulus that begins expanding at 30°/s after a 1 s static period. Frame rate: 60 fps.

**Supplementary Video S4. Isoluminant expansion combined with gray-to-black luminance change**. Video shows isoluminant annulus that begins expanding at 30°/s after a 1 s static period. A gray-to-black luminance change occurs at the annulus center 250 ms before expansion onset. Frame rate: 60 fps.

**Supplementary Video S5. Isoluminant expansion combined with gray-to-black luminance change**. Video shows isoluminant annulus that begins expanding at 30°/s after a 1 s static period. A gray-to-black luminance change occurs at the annulus center 250 ms after expansion onset. Frame rate: 60 fps.

**Supplementary Video S6. Isoluminant expansion combined with gray-to-white luminance change**. Video shows isoluminant annulus that begins expanding at 30°/s after a 1 s static period. A gray-to-white luminance change occurs at the annulus center 250 ms before expansion onset. Frame rate: 60 fps.

**Supplementary Video S7. Isoluminant expansion combined with gray-to-white luminance change**. Video shows isoluminant annulus that begins expanding at 30°/s after a 1 s static period. A gray-to-white luminance change occurs at the annulus center 250 ms after expansion onset. Frame rate: 60 fps.

